# Comparative analysis of non structural protein 1 of SARS-CoV2 with SARS-CoV1 and MERS-CoV: An i*n silico* study

**DOI:** 10.1101/2020.06.09.142570

**Authors:** Ankur Chaudhuri

## Abstract

The recently emerged SARS-CoV2 caused a major pandemic of coronavirus disease (COVID-19). Non structural protein 1 (nsp1) is found in all beta coronavirus that causes several severe respiratory diseases. This protein is considered as a virulence factor and has an important role in pathogenesis. This study aims to elucidate the structural conformations of non structural protein 1 (nsp1), prediction of epitope sites and identification of important residues for targeted therapy against COVID-19. In this study, molecular modelling coupled with molecular dynamics simulations were performed to analyse the conformational change of nsp1 of SARS-CoV1, SARS-CoV2 and MERS-CoV at molecular level. Principal component analysis escorted by free energy landscape revealed that SARS-CoV2 nsp1 protein shows greater flexibility, compared to SARS-CoV1 and MERS-CoV nsp1. From the sequence alignment, it was observed that 28 mutations are present in SERS-CoV2 nsp1 protein compared to SERS-CoV1 nsp1. Several B-cell and T-cell epitopes were identified by immunoinformatics approach. SARS-CoV2 nsp1 protein binds with the interface region of the palm and finger domain of POLA1 by using hydrogen bond and salt bridge interactions. These findings can be used to develop therapeutics specific against COVID-19.

## 1. Introduction

In December 2019, the first epidemic novel coronavirus (SARS-CoV2) was identified in Wuhan city, China [1, 2]. Since the outbreak, WHO (World Health Organization) declared the COVID-19 as a pandemic on March 11, 2020 and posed a global health emergency. The causative agent of the COVID-19 disease is a severe acute respiratory syndrome coronavirus 2 (SARS-CoV2). The worldwide number of coronavirus cases reached 27,354,078 with a death toll of 894,005 as of Sep 07, 2020 (Last accessed May 25, 2020) [3]. Although it was first reported from China, the number of active cases in India, USA, Brazil, Russia, Spain, Italy, France, Germany, and UK have surpassed the cases identified in China (85,134) (Last accessed May 25, 2020) [3]. Almost all countries initiated social distancing and locked down precautions to avoid human to human transmission for till date no human vaccine is available in the market for COVID-19 treatment. Coronaviruses are enveloped, positive-sense, single-stranded RNA viruses (ssRNA+) belonging to the Coronaviridae family. COVID-19 is a member of beta coronaviruses, like the other human coronaviruses SARS-CoV and MERS-CoV [2,4]. There are seven strains of human CoVs, which include NL63, 229E, HKU1, OC43, Middle East respiratory syndrome (MERS-CoV), severe acute respiratory syndrome (SARS-CoV or SARS-CoV1), and 2019-novel coronavirus (SARS-CoV2), responsible for the infection of the respiratory tract. Among these seven strains, three strains are highly pathogenic (SARS-CoV1, SARS-CoV2 and MERS-CoV) and are responsible for lower respiratory ailment like bronchiolitis, bronchitis, and pneumonia [5]. Sequence analysis of SARS-CoV2 suggests that the genome size of this virus is 30kb and encodes structural and non-structural proteins like other CoVs. The two-thirds of the 5′ end of the CoV genome consists of two overlapping open reading frames (ORFs 1a and 1b) that encode non-structural proteins (nsps). The other one-third of the genome consists of ORFs that encodes structural proteins. Structural protein consists of S protein (Spike), E protein (Envelope), M protein (Membrane), and N protein (Nucleocapsid) [6, 7]. The ORF1a and ORF1b encodes a polyprotein which is cleaved into sixteen non-structural proteins (nsp1-16) that form the replicase / transcriptase complex (RTC) [8]. Alpha and beta CoVs consists of 16 nsps, while gamma and delta CoVs, lacking nsp1, consists of 15 nsps (nsp2-16) [9]. The amino acid sequences of nsp1 are highly divergent among CoVs [10]. It is among the least well-understood nsps, and other than in coronaviruses, no viral or cellular homologs are reported. The nsp1 of SARS-CoV inhibits host gene expression by blocking the translation process through interaction with 40S ribosomal subunit and degrades host mRNA via the recruitment of unidentified host nuclease(s) [11, 12]. SARS-CoV nsp1 inhibits the expression of the IFN genes and the host antiviral signaling pathway in the infected cells. The dysregulation of IFN genes is the key factor for inducing lethal pneumonia [13, 14]. MERS-CoV nsp1 also induced mRNA degradation and translational suppression. SARS-CoV nsp1 also regulates the induction of cytokines and chemokine in human lung epithelial cells [15]. Thus, nsp1 is considered a major possible virulence factor for coronaviruses. SARS-CoV2 nsp1 antagonizes interferon induction to suppress the host antiviral response. The inflammatory phenotype of SARS-CoV and SARS-CoV2 pathology was also contributed by nsp1 protein [15]. The nsp1 protein of SARS-CoV2 interacts with six proteins of the infected host cells. They are POLA1, POLA2, PRIM1, PRIM2, PKP2 and COLGALT1. Four of these host proteins (POLA1, POLA2, PRIM1 and PRIM2) form DNA polymerase alpha complexes. These events raise the possibility that the nsp1 protein of SARS-CoV2 may interact with the DNA polymerase alpha complex and change its functional activity to antagonise the innate immune system [8].

The main goal of this study is to propose the molecular model of the three nsp1 proteins and SARS-CoV2 nsp1-POLA1 complex. The epitope sites of SARS-CoV1, SARS-CoV2 and MERS-CoV nsp1 protein were identified by immunoinformatics process. Molecular dynamics simulation, principal component analysis (PCA), and Gibbs free energy landscape (FEL) were performed to evaluate the structural flexibility and dynamic stability of the SARS-CoV1, SARS-CoV2 and MERS-CoV nsp1 protein. The mutated amino acids of SARS-CoV2 nsp1 protein were reported by using multiple sequence alignment. The binding interactions of SARS-CoV2 nsp1 protein with its host cell receptor POLA1 were studied by implementing protein-protein docking procedure to identify the important interacting residues at the interface region. The all-atom MD simulations for 50 ns on the three nsp1 proteins were performed to describe conformational changes under explicit solvent conditions. This study delivers an atomistic insight into the structure and conformation of nsp1 of SARS-CoV1, SARS-CoV2 and MERS-CoV by utilizing state-of-the-art computational approaches.

## 2. Materials and methods

### 2.1 Sequence retrieval

The protein sequences of nsp1 were retrieved from the curated NCBI database [16]. The accession numbers of the nsp1 of SARS-CoV1, SARS-CoV2 and MERS-CoV are NP_828860.2, YP_009725297.1 and YP_009047213.1 respectively. The pairwise sequence identity between COVID-19 nsp1 protein and each of the other HCoV nsp1 proteins (SERS-CoV1 and MERS-CoV) was calculated using the BLASTp (basic local alignment tool) [17]. To check the conservation pattern, multiple sequence alignment (MSA) of all of the nsp1 sequences was performed using the Clustal Omega programme of the European Bioinformatics Institute (EMBL-EBI) [18].

### 2.2 Three dimensional structure prediction

The NMR structure of the non structural protein 1 (nsp1) of SERS-CoV1 (residues 13-127) was identified by Almeida et al. 2007 and deposited in Protein Data Bank (PDB) as ID no. 2hsx/2gdt [19]. Currently no crystallographic structure of nsp1 of SARS-CoV2 and MERS-CoV is available in Protein Data Bank (PDB). So, in silico modelling study was employed to predict the three dimensional structure of nsp1 of SARS-CoV2 and MERS-CoV by using the I-TASSER web server [20]. I-TASSER (Iterative Threading ASSEmbly Refinement) is a bioinformatics approach to predict the structure and function of an unknown protein molecule. It first detects structural templates from the protein data bank database by fold recognition or multiple threading approach LOMETS [21]. The full-length atomic models are constructed by iterative template-based fragment assembly simulations. The predicted protein models are constructed by continuous assembling of the aligned region with templates and initiating an *ab initio* folding for the unaligned regions based on replica exchange Monte Carlo simulation process. This simulation method generates an ensemble of several conformations which are further clustered on the basis of free energy. The lowest energy predicted structures are subjected to a refinement process resulting in a final three dimensional protein structural model. I-TASSER (as ‘Zhang-Server’) has regularly been the top ranked server for prediction of protein structure in recent community-wide CASP (Critical Assessment of Protein Structure Prediction) method experiments [22]. The modelled nsp1 proteins were optimised to avoid any stereochemical restraints by steepest descent energy minimization method. The stereochemical quality of the nsp1 proteins was validated by Ramachandran plot using PROCHECK [23, 24]. The models were further validated by ProSA [25], ProQ [26].

### 2.3 Active site prediction

The ligand binding residues of nsp1 of SARS-CoV1, SARS-CoV2 and MERS-CoV were predicted by COACH meta server [27]. COACH generates complementary ligand binding sites of the target proteins by using two comparative processes, S-SITE and TM-SITE. These two methods recognize ligand-binding templates from the BioLiP database [28] by sequence profile comparisons and binding-specific substructure.

### 2.4 Prediction of T-cell (HLA class I and II) epitopes

The RANKPEP, sequence-based screening server was used to identify the T-cell epitopes [29] of the nsp1 protein of SARS-CoV1, SARS-CoV2 and MERS-CoV. This server predicts the short peptide that binds to MHC molecules from protein sequences using the position-specific scoring matrix (PSSM). All the HLA class I alleles were selected for prediction of epitopes of HLA class I. For the prediction of epitopes of HLA class II, we selected some alleles such as DRB10101, DRB10301, DRB10401, DRB10701, DRB10801, DRB11101, DRB11301, and DRB11501 that cover HLA variability of over 95% of the human population worldwide [30].

### 2.5 B-cell epitopes (linear) identification

B-cell epitopes of the three nsp1 proteins were predicted by using BepiPred and Kolaskar & Tongaonkar Antigenicity (http://www.iedb.org/) servers [31]. BepiPred for linear epitope prediction uses both amino acid propensity scales and hidden Markov model methods. The cut off score for linear B-cell epitopes prediction is 0.50. Kolaskar and Tongaonkar evaluate the protein for B cell epitopes using the physicochemical properties of the amino acids and their frequencies of occurrence in recognized B cell epitopes [32, 33].

### 2.6 Molecular dynamics simulation

Molecular dynamics simulations are used to predict the dynamic behaviour of the protein macromolecule at the atomic level. The nsp1 protein of SERS-CoV1, SARS-CoV2 and MERS-CoV were subjected to MD simulation by using Gromacs v 2018.2 software suite [34] with OPLS-AA force field [35]. The three systems were solvated in a cubic box with SPC (simple point charge) water model [36] by maintaining periodic boundary condition (PBC) through the simulation process. Sodium and chloride ions were added to neutralize the three systems. Each system was energy minimized using the steepest descent algorithm until the maximum force was found to be smaller than 1000.0 kJ/mol/nm. This was done to remove any steric clashes on the system. Each system was equilibrated with 100 ps isothermal-isochoric ensemble, NVT (constant number of particles, volume, and temperature) followed by 100 ps isothermal-isobaric ensemble NPT (constant number of particles, pressure, and temperature). The two types of ensemble of equilibration method stabilized the three systems at 310 K and 1 bar pressure. The Berendsen thermostat and Parrinello-Rahman were applied for temperature and pressure coupling methods respectively [37]. Particle Mesh Ewald (PME) method [38] was used for the calculations of the long range electrostatic interactions and the cut off radii for Van der Waals and coulombic short-range interactions were set to 0.9 nm. The Linear Constraint Solver (LINCS) constraints algorithm was used to fix the lengths of the peptide bonds and angles [39]. All the three systems were subjected to MD simulations for 50 ns. The resulting MD trajectories were utilized through the inbuilt tools of GROMACS for analysis purposes. The subsequent analyses were performed using VMD [40], USCF Chimera [41], Pymol [42], and also the plots were created using xmgrace [43].

### 2.7 Principal components analysis (PCA) or essential dynamics

Principal components analysis or essential dynamics is a process which extracts the essential motions from the MD trajectory of the targeted protein molecule [44]. The nsp1 protein of SARS-CoV1, SARS-CoV2 and MERS-CoV are used for this purpose. The initial step of PCA analysis is to construct the covariance matrix which examines the linear relationship of atomic fluctuations for individual atomic pairs. The diagonalization of covariance matrix results in a matrix of eigenvectors and eigenvalues. The eigenvectors determine the movement of atoms having corresponding eigenvalues which represents the energetic contribution of an atom participating in motion. The covariance matrix and eigenvectors were analyzed using the *gmx cover* and *gmx anaeig* tool respectively. Gibbs free-energy landscape (FEL) elaborates the protein dynamic processes by representing the conformational states and the energy barriers [45]. The FEL of SARS-CoV1, SARS-CoV2 and MERS-CoV was constructed based on the first (PC1) and second (PC2) principal components, Rg and rmsd, and psi and phi angels. FEL was calculated and plotted by using *gmx sham* and *gmx xpm2ps* module of GROMACS.

### 2.8 Protein-protein docking

Nsp1 protein of SARS-CoV2 interacts with POLA1 and blocks the host cell replication process. [8]. The molecular interactions of nsp1 protein of SERS-CoV2 (COVID-19) with the catalytic subunit of human DNA polymerase alpha, POLA1 was analyzed by using the HADDOCK (High Ambiguity Driven protein-protein DOCKing) programme. It is a flexible docking approach for the modelling of biomolecular complexes. It encodes instruction from predicted or identified protein interfaces in ambiguous interaction restraints (AIRs) to drive the docking procedure [46]. The coordinates of the solved structure of the catalytic domain of DNA polymerase alpha, POLA1 was downloaded from PDB database (PDB ID: 6AS7) and prepared for the docking experiments by removing water, ions and the ligands. The interface residues were utilized for the docking procedure. The active residues of POLA1 (Asp860, Ser863, Leu864, Arg922, Lys926, Lys950, Asn954 and Asp1004) were retrieved from the literature [47]. The active residues (Leu16, Leu18, Phe31, Val35, Glu36, Leu39, Arg43, Leu46, Gly49, Iso71, Pro109, Arg119, Val121 and Leu123) of nsp1 of SARS-CoV2 were predicted from COACH server. The amino acids surrounding the active residues of both proteins were selected as passive in docking procedure. Active residues are the amino acids from the interface region of the two proteins that take part in direct binding with the other protein partner while passive residues are the amino acids that can interact indirectly in docking procedure. Approximately 163 structures in 8 clusters were obtained from HADDOCK server, which represented 81.5% of the water-refined models. PRODIGY software [48] was used to predict the binding affinity and dissociation constant for each SARS-CoV2 nsp1-POLA1 complex from the best three clusters. The generated model of SARS-CoV2 nsp1-POLA1 complex was optimized to avoid any stereochemical restraints by steepest descent energy minimization method. The optimized complex was validated by Ramachandran plot analysis of ψ/φ angle from PROCHECK. PISA server (http://www.ebi.ac.uk/msd-srv/prot_int/, Last accessed May 25, 2020) was used to calculate total buried surface area, nature of interactions and amino acids involved in interactions at interface region.

## 3. Results and discussion

### 3.1 Sequence analysis and protein structure prediction

The sequence identity of nsp1 protein of SARS-CoV2 with SARS-CoV1 and MERS-CoV was 84.4% and 20.61% respectively. Multiple sequence alignment (MSA) of nsp1 proteins was performed to identify the conserved residues. The amino acids marked as asterisk illustrate the positions of nsp1 protein that were highly conserved over the evolutionary time scale (**Fig. 1**). The differences in the amino acid changes were also recorded. It was observed that compared to SARS-CoV1, there were 28 mutations in the nsp1 protein of SARS-CoV2 (**Fig. 1**). The three dimensional structure of nsp1 of SARS-CoV2 and MERS-CoV was modelled using the I-TASSER web server (**Fig. 2**). The NMR structure of SARS-CoV1 nsp1^13–127^ (PDB: 2hsx/2gdt) was identified as the template for three dimensional structure prediction of SARS-CoV2 nsp1 on the I-TASSER server. The highly significant templates used in the modelling of the nsp1 protein of SARS-CoV2 and MERS-CoV through I-TASSER are listed in **Table 1**. It was evident from the high Z score (>1 means good alignment) and good coverage in case of most of the structural templates, the generated threading alignment predicts a good and confident model in both cases. As shown from the topological analysis, SARS-CoV2 nsp1 ^13–127^ exhibits similar α/β-folds with SARS-CoV nsp1^13–127^ that consist of a characteristic six β-barrel and a long α-helix. It was also observed that a short α helix (163-168) is present in the C-terminal region of the of the SARS-CoV2 nsp1 protein. This α-helix may play an important role in the inhibition of the host protein synthesis process [49]. In case of MERS-CoV, eight β-barrel and four α-helix constitutes the nsp1 protein which is different from the SARS-CoV1 and SARS-CoV2 (**Fig. S1**). The stereochemical quality of the model nsp1 proteins was validated on the basis of Ramachandran analysis of ψ/φ angle from PROCHECK. Examination of Ramachandran plot of nsp1 protein of SARS-CoV2 and MERS-CoV showed above 92% residues lie in the allowed regions (**Table S1**). From ProSA and ProQ analysis, it is clear that the overall model quality of the nsp1 protein of SARS-CoV2 and MERS-CoV are within the range of scores typically found for proteins of similar size (**Table S1)**. The important residues of the nsp1 protein of SARS-CoV1, SARS-CoV2 and MERS-CoV that are involved in the ligand binding process are listed in **Table 2**. Combination of the finding of the different domains of SARS-CoV1, SARS-CoV2, and MERS-CoV may help better understanding of the entire structure of nsp1.

**Table 1:** The highly significant structural templates for sequence alignment obtained from PDB library for modelling through I-TASSER

| **SARS-CoV2** |  | **MERS-CoV** |  |
| --- | --- | --- | --- |
| PDB ID | Normalized Z Score | PDB ID | Normalized Z Score |
| 2hsxA | 1.83 | 2hsx | 8.11 |
| 2gdtA | 2.52 | 2hsx | 9.76 |
| 2gdt | 1.70 | 2hsx | 5.19 |
| 2hsx | 10.03 | 5xbc | 1.43 |
| 2hsx | 4.59 | 3zbd | 1.14 |
| 2hsxA | 2.14 | 2v0gA | 0.72 |
| 2hsx | 7.61 | 2gdtA | 0.95 |
| 2gdtA | 3.07 | 2p97A | 0.63 |
| 2gdtA | 2.75 | 1v8eA | 0.60 |
| 2hsxA | 2.34 | 3fdfA | 0.69 |

**Table 2:** Active residues prediction by COACH server

| **Nsp1 proteins** | **Active site residues** |
| --- | --- |
| SARS-CoV1 | Leu16, Leu18, Phe31, Leu39, Arg43, Leu46, Gly49, Iso71, Pro109, Arg119, Val121, Leu123 |
| SARS-CoV2 | Leu16, Leu18, Phe31, Val35, Glu36, Leu39, Arg43, Leu46, Gly49, Iso71, Pro109, Arg119, Val121, Leu123 |
| MERS-CoV | Arg13 |

**Fig. 1:**
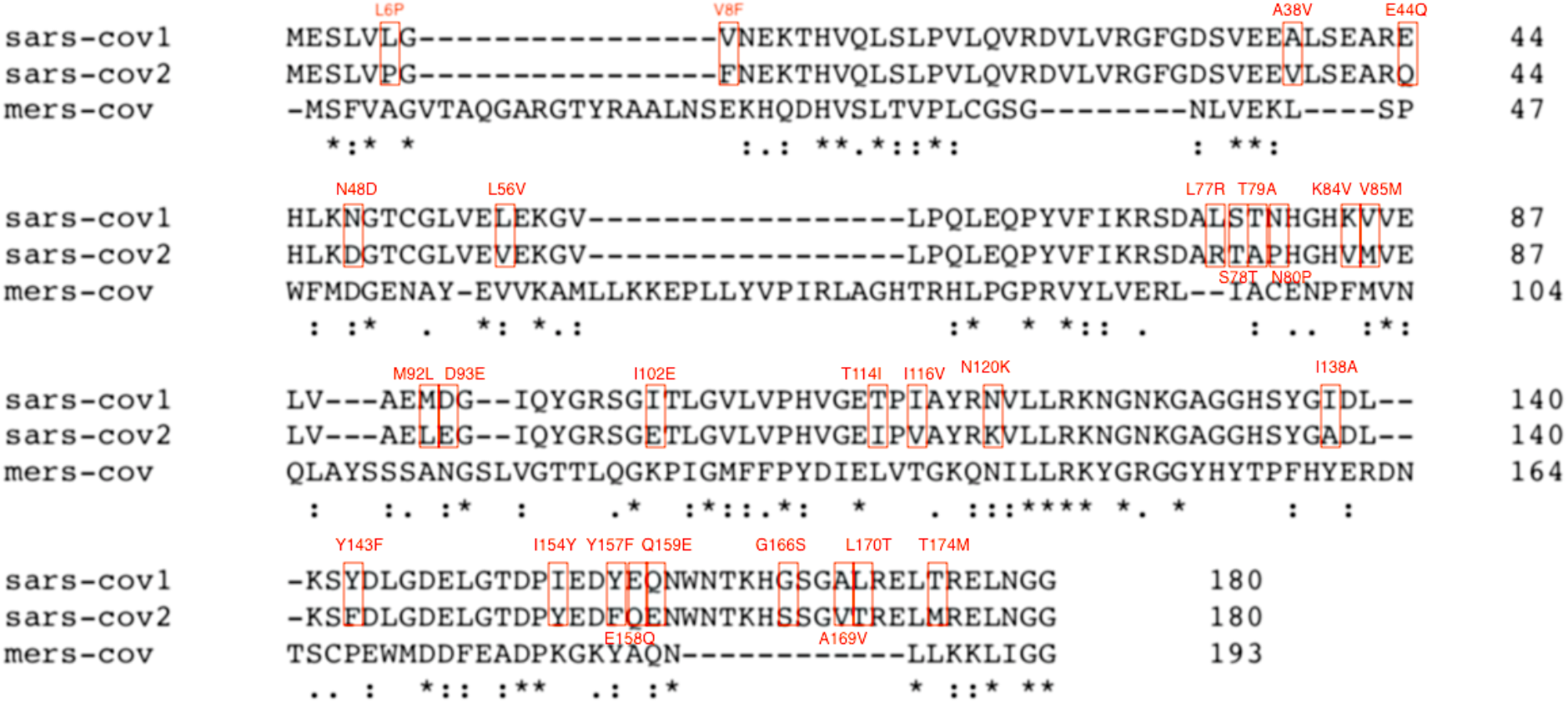
Multiple sequence alignment for nsp1 protein of SARS-CoV1, SARS-CoV2 and MERS-CoV. The alignment is shown using the Clustal Omega web server. Asterisk represents the conserved residues. Mutated residues of SARS-CoV2 nsp1 protein are represented as red box

**Fig. 2:**
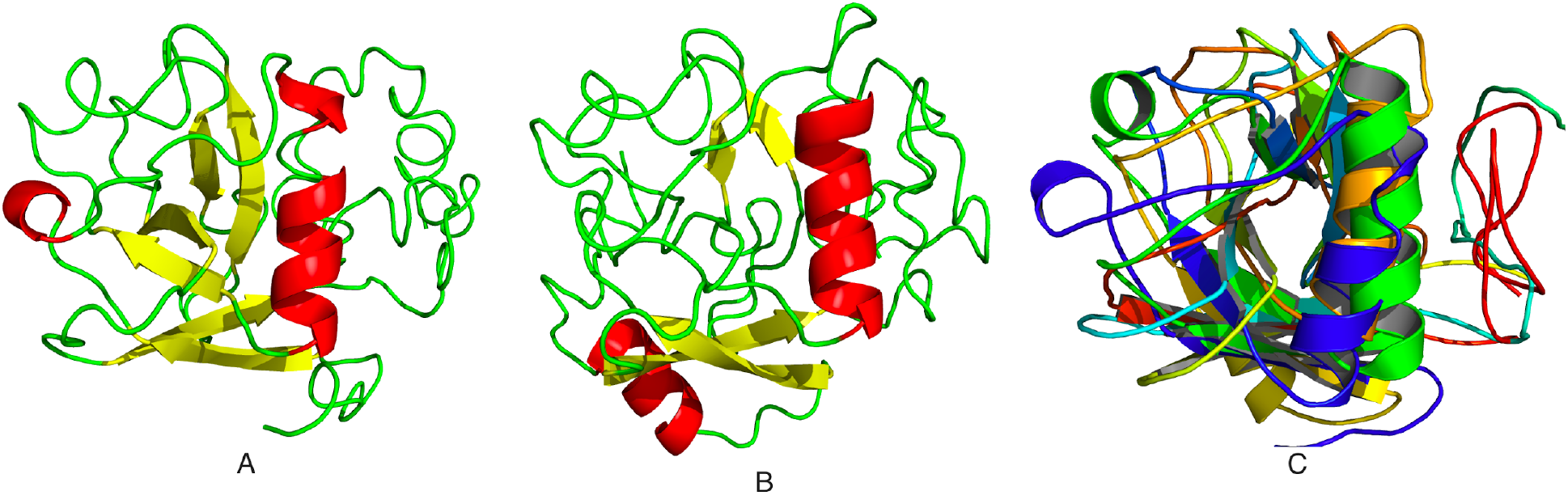
Prediction of three dimensional structure of nsp1 protein by I-TASSER programme. A. Modelled structure of SARS-CoV2 nsp1 protein. B. Modelled structure of MERS-CoV nsp1 protein. C. Superimposition of SARS-CoV1 (PDB ID: 2hsx) with SARS-CoV2 and MERS-CoV nsp1 (Orange, blue and green colour represents the SARS-CoV1, SARS-CoV2 and MERS-CoV nsp1, respectively)

### 3.2 Defining T-cell and linear B-cell epitopes

Several studies revealed that specific T-cell response is required for the elimination of several viral infections such as influenza A, SARS-CoV, MERS-CoV and para-influenza. These studies conclude that T-cell mediated response is essential for the development of specific vaccines [50, 51]. CD8^+^ cytotoxic T-cells recognize the infected cells in the lungs whereas CD4^+^ helper T-cells are essential for the production of specific antibodies against viruses. Here we used RankPep server to predict peptide binders to MHC class I and MHC class II alleles from nsp1 protein sequences by using Position Specific Scoring Matrices (PSSMs). The antigenic epitopes of three nsp1 proteins with high binding affinity were predicted and summarized in **Table 3 and Table 4**. Secreted neutralising antibodies play an important role to protect the body against viruses. The entry process of the viruses is blocked by the SARS-CoV specific neutralizing antibodies [52]. The Bepipred web server was employed for the linear B-cell epitope prediction study. SARS-CoV1, SARS-CoV2 and MERS-CoV nsp1 proteins were used for this purpose. The Kolaskar & Tongaonkar Antigenicity method was employed for the cross-checking of the predicted epitopes. The linear B-cell epitopes of the three nsp1 proteins are depicted in **Table 5**. Both humoral and cellular immune responses are important factors against coronavirus infection [52]. Finally, in SARS-CoV2 nsp1 protein, four epitope rich regions (15-27, 45-81, 121-140 and 147-178) that were shared between T-cell and B-cell were reported. This information will be helpful for vaccine design by targeting SARS-CoV2 nsp1 protein.

**Table 3:**
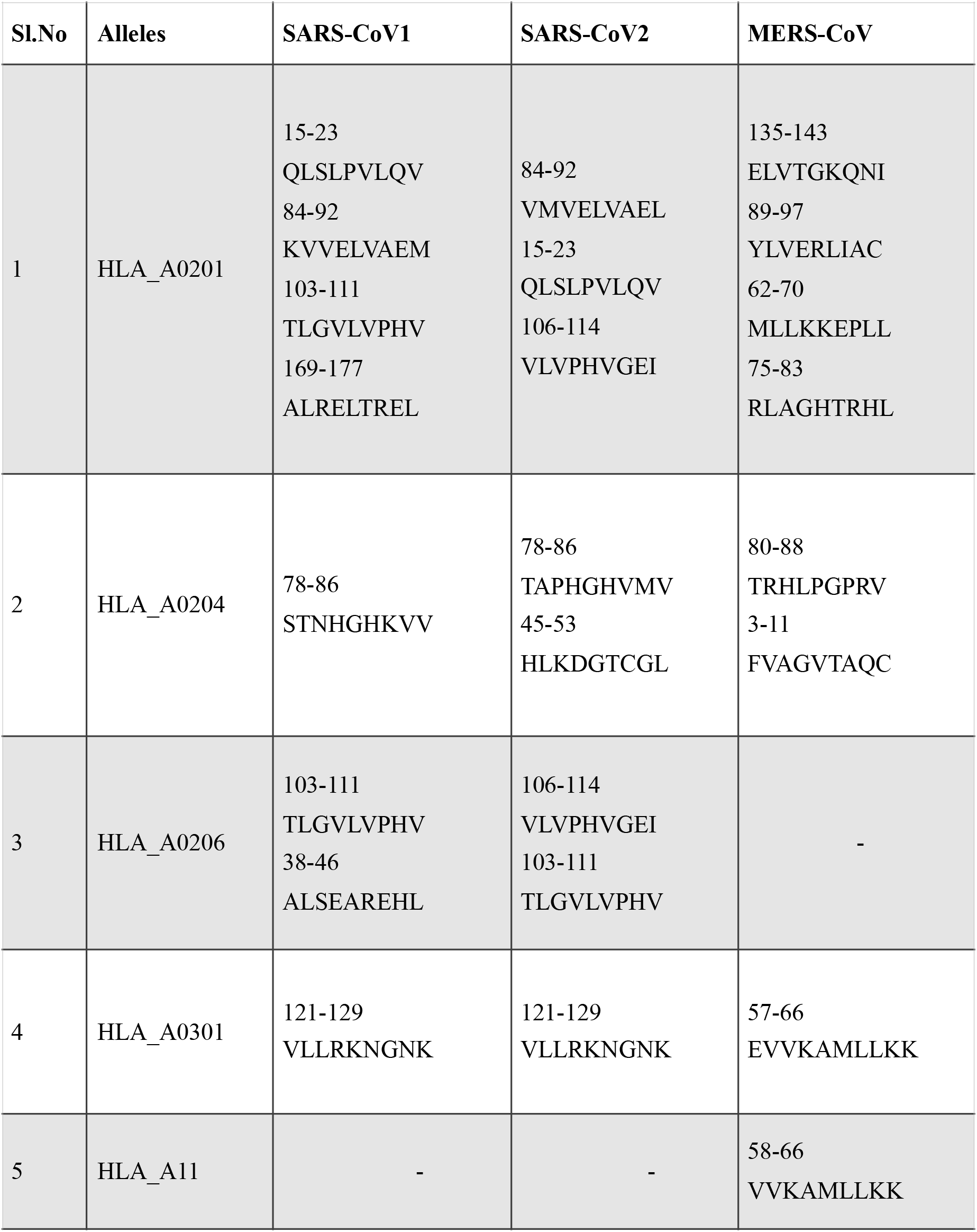
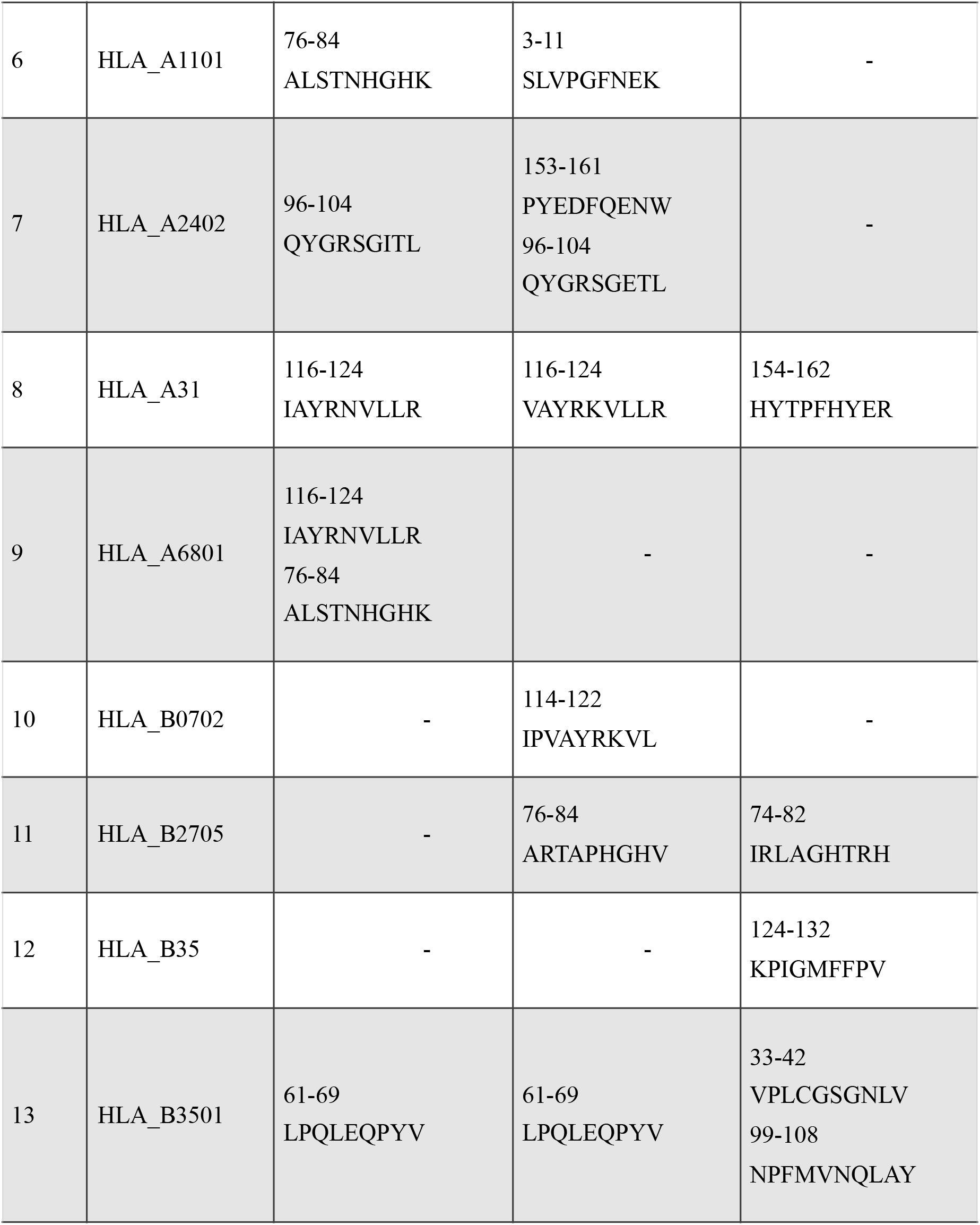
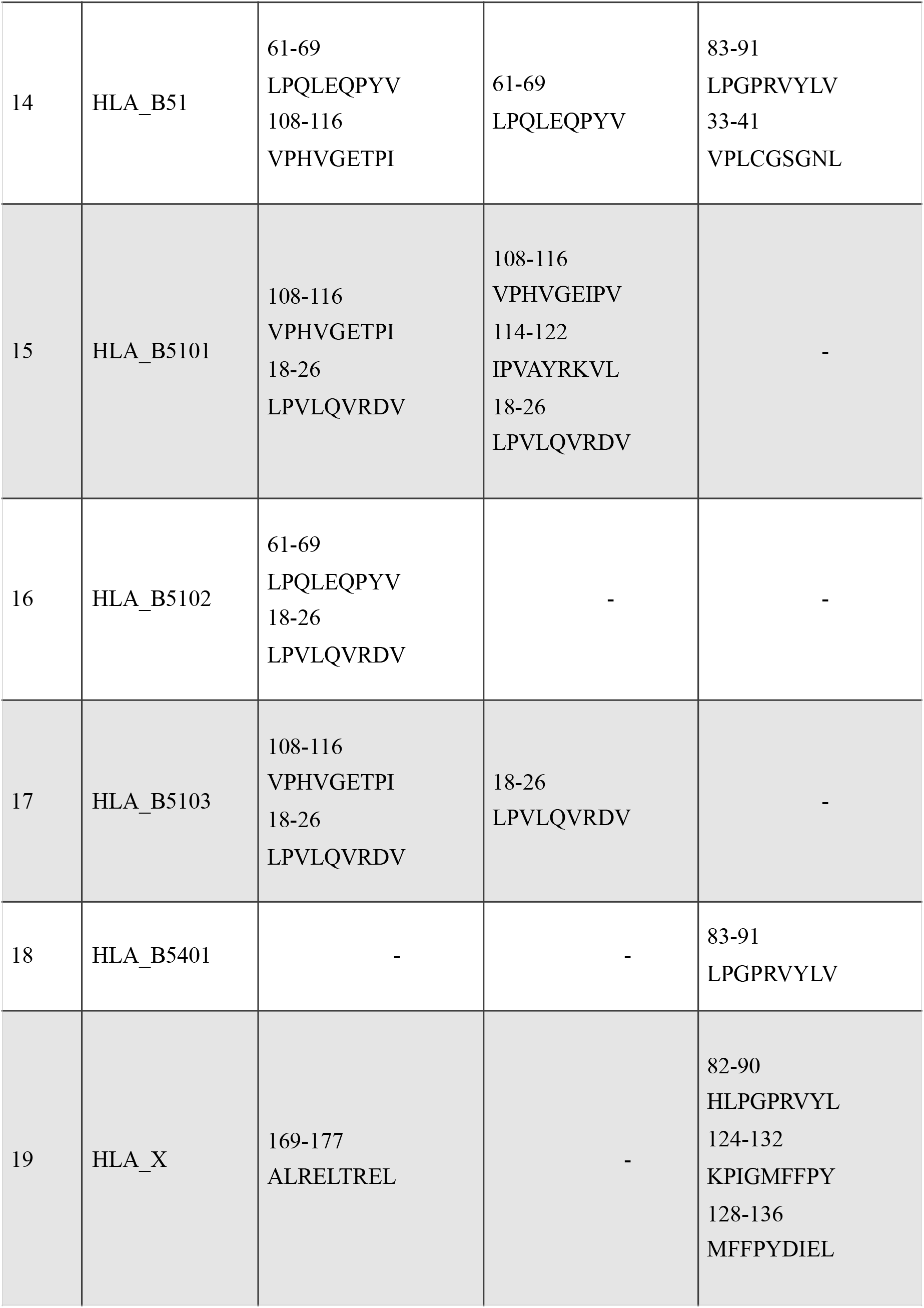
HLA I antigenic epitopes predicted using Rankpep

**Table 4:**
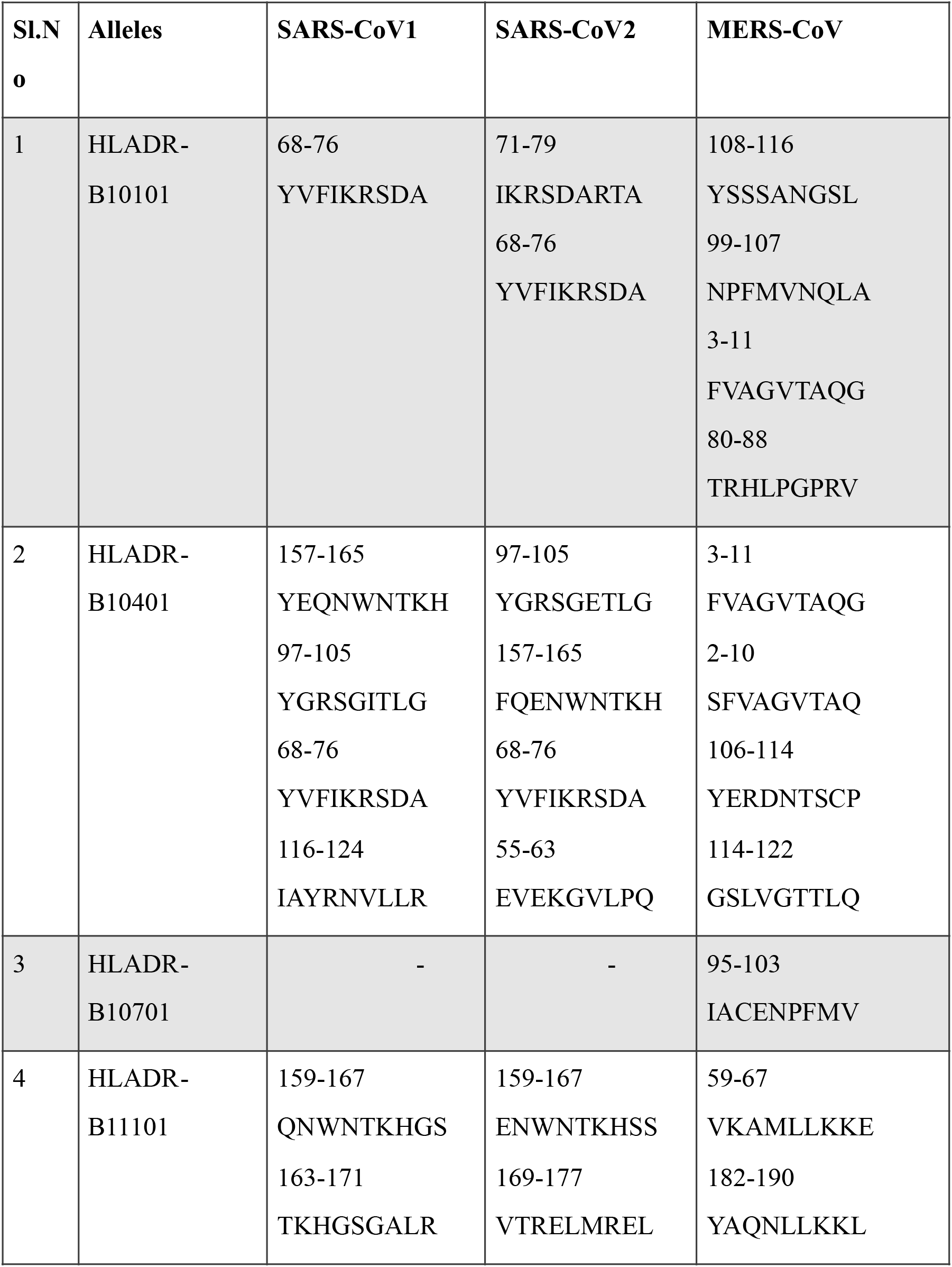
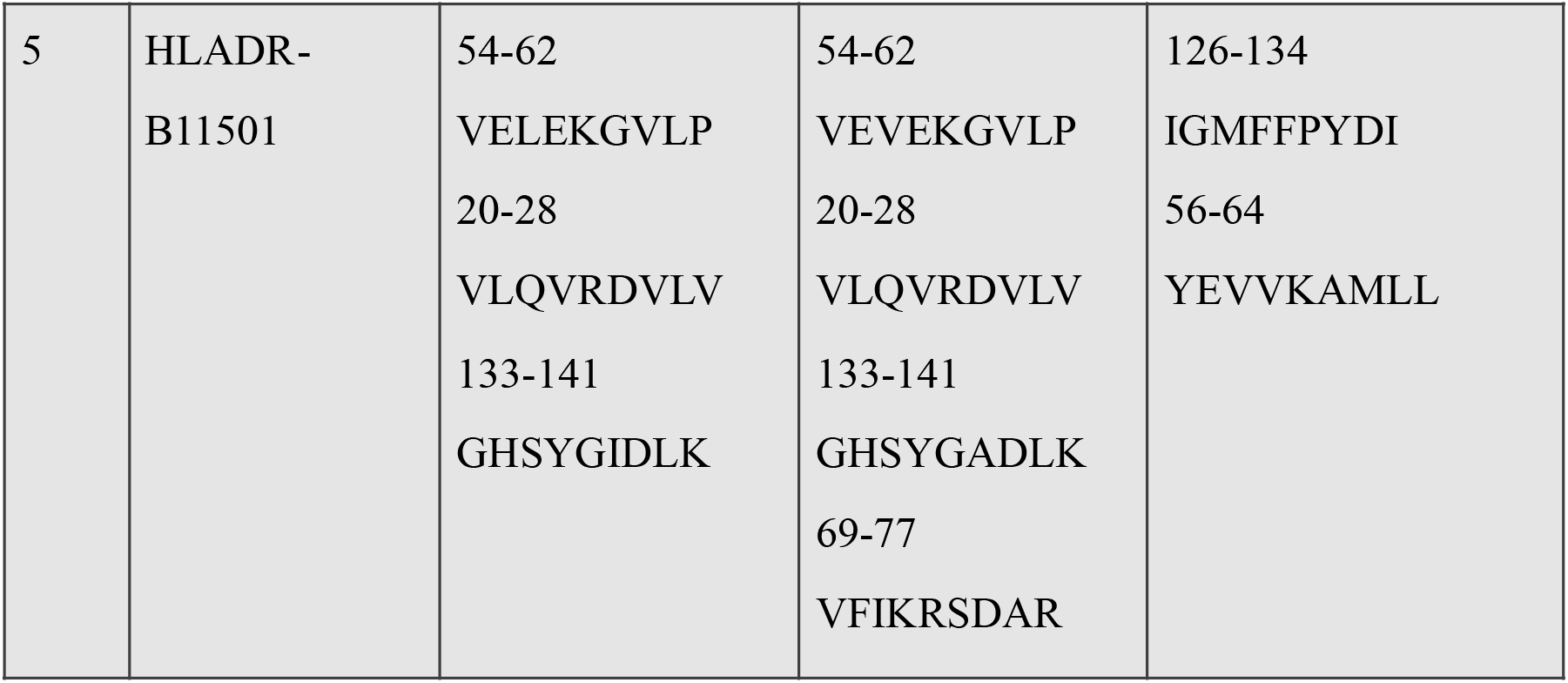
HLA II antigenic epitopes predicted using Rankpep

**Table 5:**
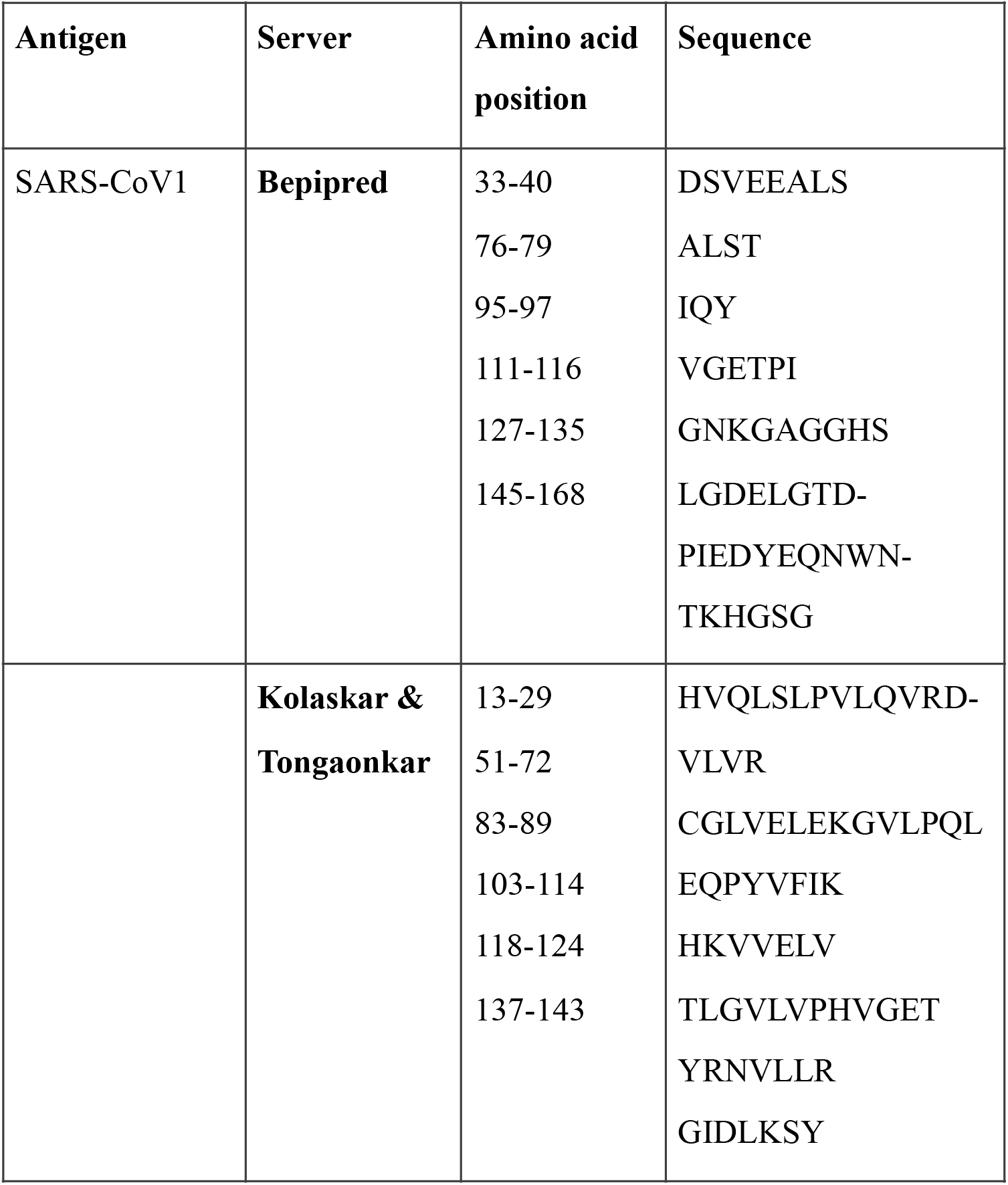
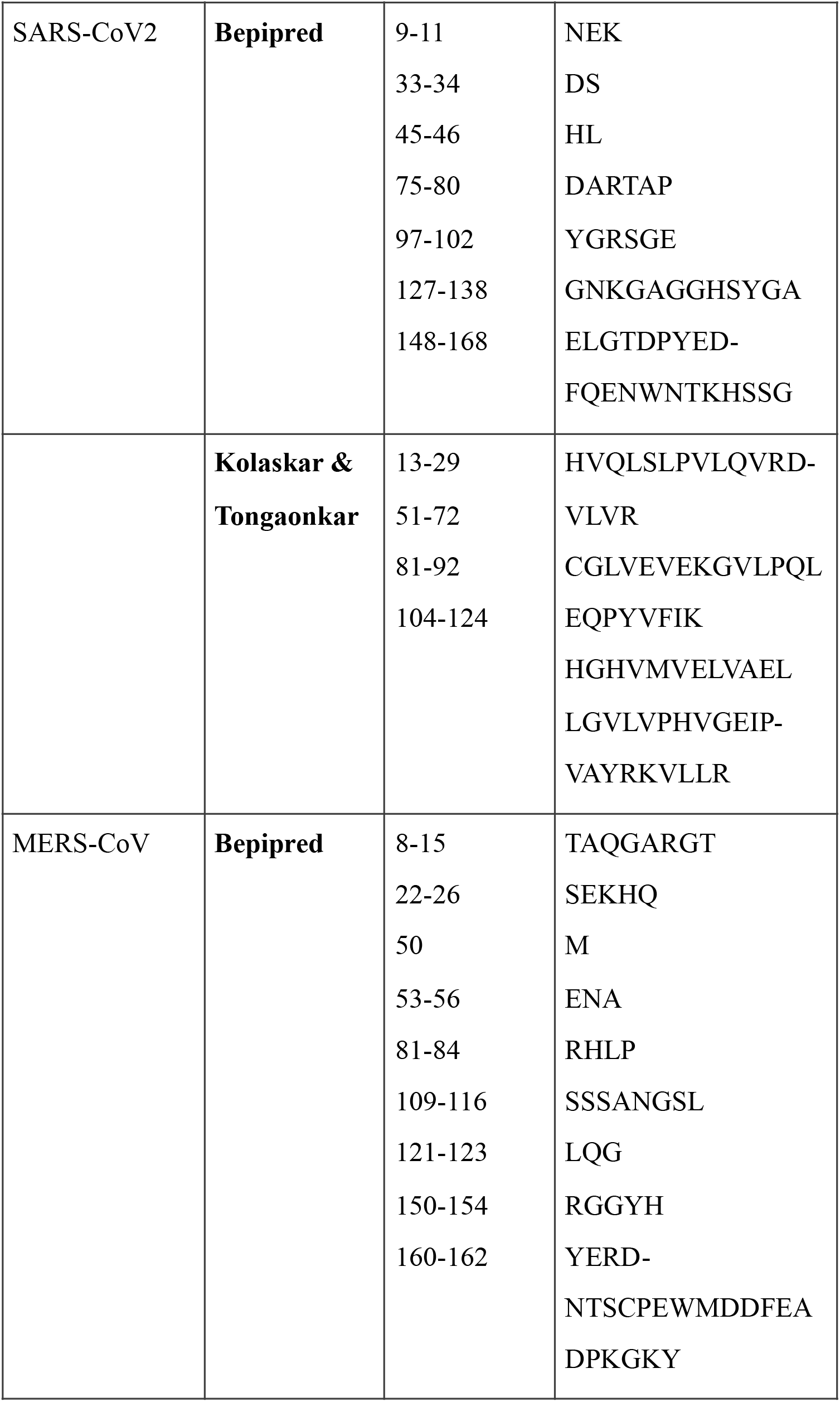
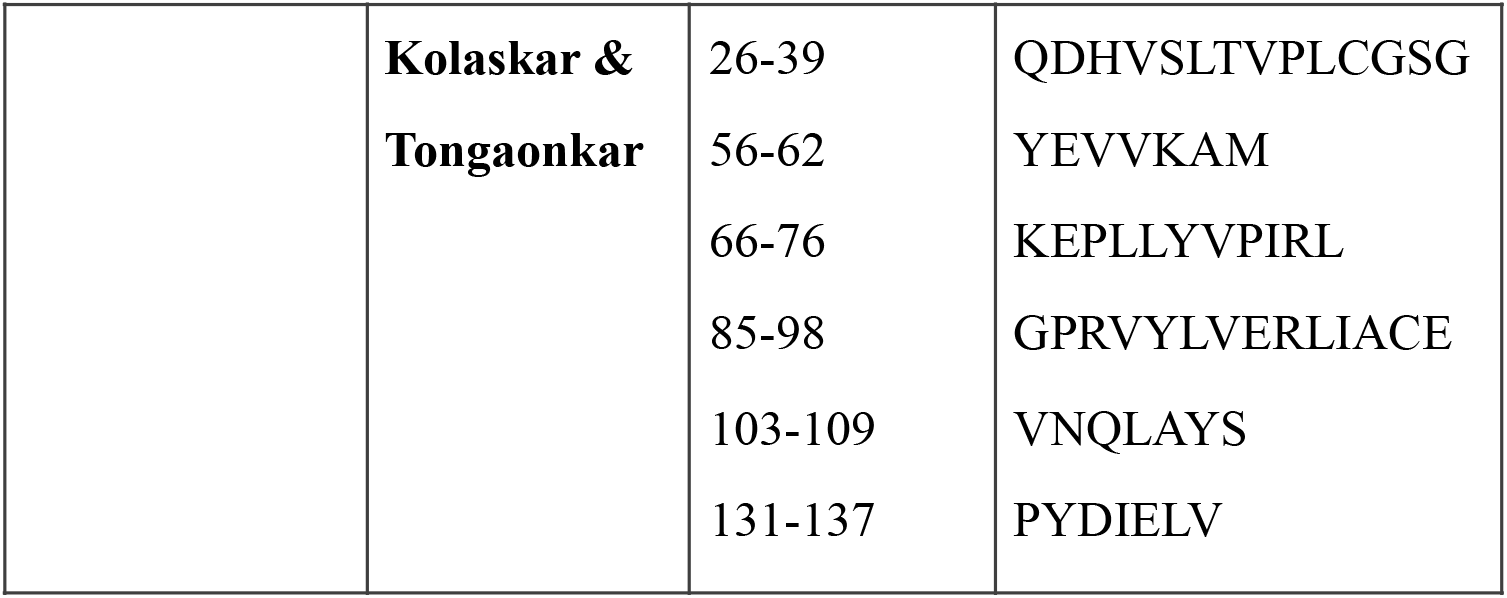
Predicted linear B-cell epitopes of SARS-CoV1, SARS-CoV2 and MERS-CoV nsp1 protein via Bepipred and Kolaskar & Tongaonkar antigenicity

### 3.3 MD simulation analysis

MD simulation methods have been utilized as an accompaniment to explore the conformational behavior of proteins. Here, to understand the structural behavior of three nsp1 proteins, we have performed MD simulation studies for 50 ns. To determine the equilibration and system stability during the simulations of three nsp1 proteins, the backbone RMSDs, radius of gyration (Rg), solvent accessible surface area (SASA), and root mean square fluctuations (RMSF) were calculated and monitored over the course of 50 ns simulations and are represented in **Figure 3**. Evaluation of the structural drift was provided by the analysis of the RMSDs from the initial structures as a function of simulation time. The resulting RMSD graph was explored to assess the conformational changes during simulation. The nsp1 of SARS-CoV1 shows a smallest deviation during the initial 0-15 ns and achieved equilibrium at ∼15 ns and it remained stable for the period of 50 ns. This suggests no significant changes observed during simulation. The backbone RMSD of SARS-CoV2 nsp1 increases sharply till 14 ns and reaches a value from 0.2 nm to 0.7 nm. Thereafter, it showed flexible stability at ∼0.7 to 0.85 nm during ∼ 18 to 44 ns. After 44 ns RMSD drop down. This maximum RMSD fluctuation indicated that the SARS-CoV2 nsp1 was undergoing a large conformational change during the simulation. In case of MERS-CoV nsp1, backbone RMSD deviates ∼ 0.44 nm during first 8 ns and remained stable thereafter. (**Fig. 3A**). The average value of backbone RMSD of SARS-CoV1, SARS-CoV2 and MERS-CoV was 0.25 nm, 0.77 nm and 0.44 nm, respectively (**Fig. 3A**). SASA analysis suggested that the exposure of the three nsp1 protein surfaces to the solvent and the changes in solvent accessibility could lead to conformational changes of the nsp1 proteins. **Fig. 3B** shows the variations in SASA for the SARS-CoV1, SARS-CoV2 and MERS-CoV nsp1 protein with respect to simulation time. The average value of SASA of SARS-CoV1, SARS-CoV2 and MERS-COV was 68 nm^2^, 110 nm^2^, and 118 nm^2^, respectively. The SASA values for the SARS-CoV1 were reduced when compared with the case of SARS-CoV2 and MERS-CoV. Overall, the SASA of three systems seems to attain a stable equilibrium without any major peak during the simulation, signifying the stability of compactness. Root mean square fluctuations (RMSFs) of each amino acid highlights the flexible regions of the three nsp1 systems. RMSFs values higher than 0.25 nm are characteristic of amino acid residues belonging to flexible regions. The RMSF plot of SARS-CoV1 nsp1 showed very less flexible sites (67, 70, 116) during the simulation. The most significant conformational shifts occurred in the seven regions namely 1-3, 6-11, 32-36, 58-60, 80-81, 99, 126-129, 130-138, 144-148, 150-160 and 165-180 of SARS-CoV2 nsp1 protein. The higher fluctuations are related to the residues of a loop and helix regions. The RMSF analysis of MERS-CoV1 nsp1 showed that residues 1, 18-21, 49-52, 98-99, 113-119, 193 exhibited fluctuations during simulation (**Fig. 3C**). The high RMSF values of SARS-CoV2 indicated a larger degree of flexibility in this protein compared with SARS-CoV1 and MERS-CoV (**Fig. 3C**). To further comprehend the structural stability of three nsp1 proteins, we studied the compactness parameter of the structure by figuring the radius of gyration (Rg). Radius of gyration (Rg) is detected as root mean square deviation between the center of gravity of the respected protein and its end. It detects the stability and firmness of the simulation system and changes over simulation time due to protein folded-unfolded states [53]. The average Rg value was 1.34 nm, 1.69 nm and 1.61 nm for SARS-CoV1, SARS-CoV2 and MERS-CoV respectively. The Rg plot of SARS-CoV2 showed a smallest deviation till 14 ns. Thereafter it showed flexible stability. However, the *R*g plot shows two small drifts during 50 ns simulation. The gyration curve showed a decrease in the overall Rg value of the SARS-CoV1 nsp1 protein compared with the SARS-CoV2 and MERS-CoV, indicating that nsp1 protein of SARS-CoV1 was in a compactly packed state and had stable folding (**Fig. 3D**). From RMSD, RMSF and Rg analysis, it was observed that SARS-CoV2 shows flexible stability during MD simulation. An overall trend of backbone RMSD, SASA, RMSF and radius of gyration indicated that all three nsp1 protein systems were well equilibrated and stable during the simulation run. Hydrogen bonds provide most of the directional interactions that represents protein structure, protein folding and molecular recognition. The core of most protein structures is composed of secondary structures such as α-helix and β-sheet, which are stabilized by H-bonds between main chain carbonyl oxygen and amide nitrogen. The formation of hydrogen bonds as a function of simulation time was analyzed. The average number of hydrogen bonds of SARS-CoV1, SARS-CoV2 and MERS-CoV were 69±5, 92±10 and 90±11, respectively **(Fig. 4)**.

**Fig. 3:**
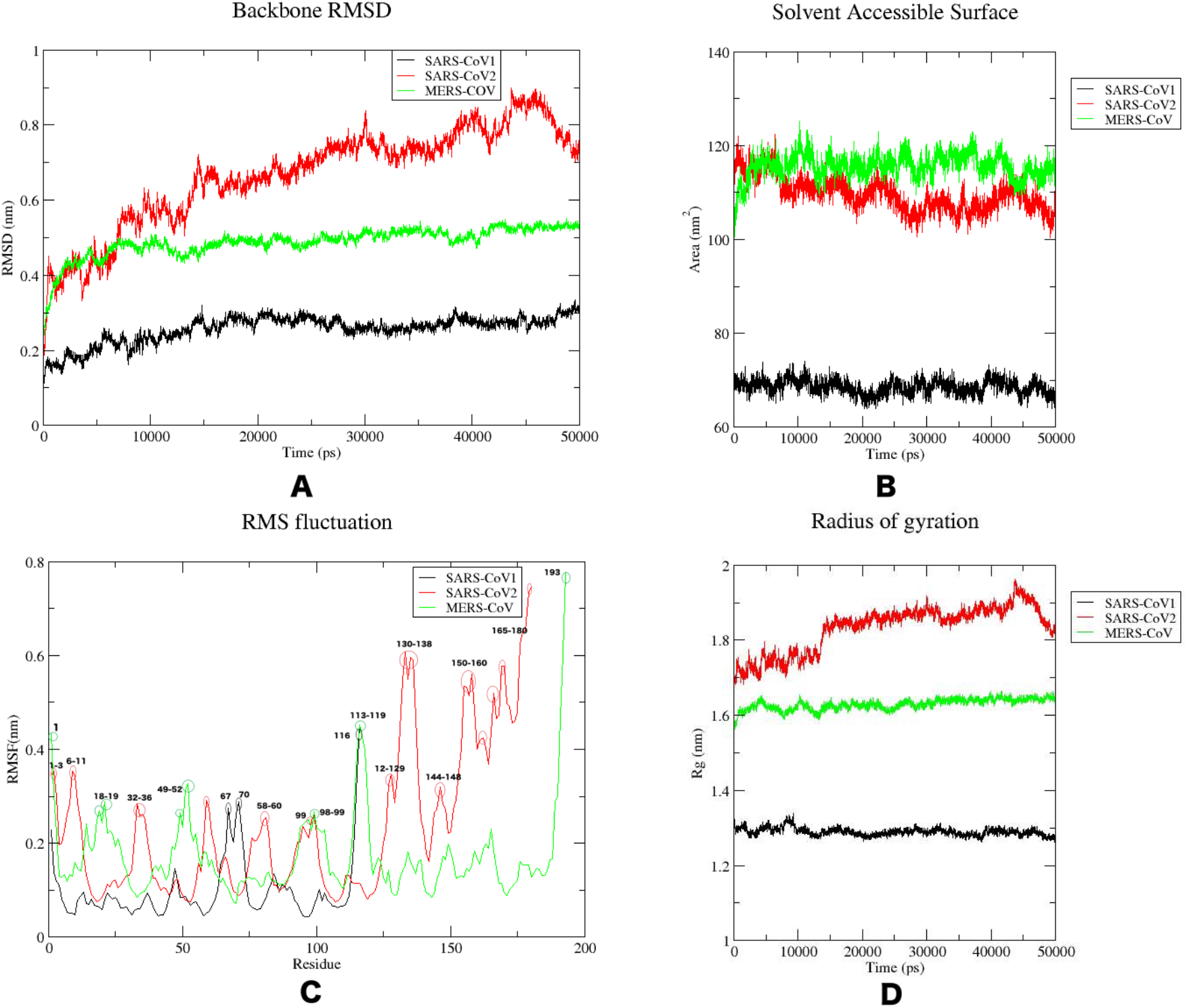
Analysis of MD simulation of three nsp1 proteins. A. Root mean square deviation (RMSD). B. Solvent accessible surface area (SASA). C. Root mean square fluctuations (RMSF). D. Radius of gyration. Black, red and green colour represents the SARS-CoV1, SARS-CoV2 and MERS-CoV nsp1, respectively. Flexible regions of each nsp1 are represented as circle

**Fig. 4:**
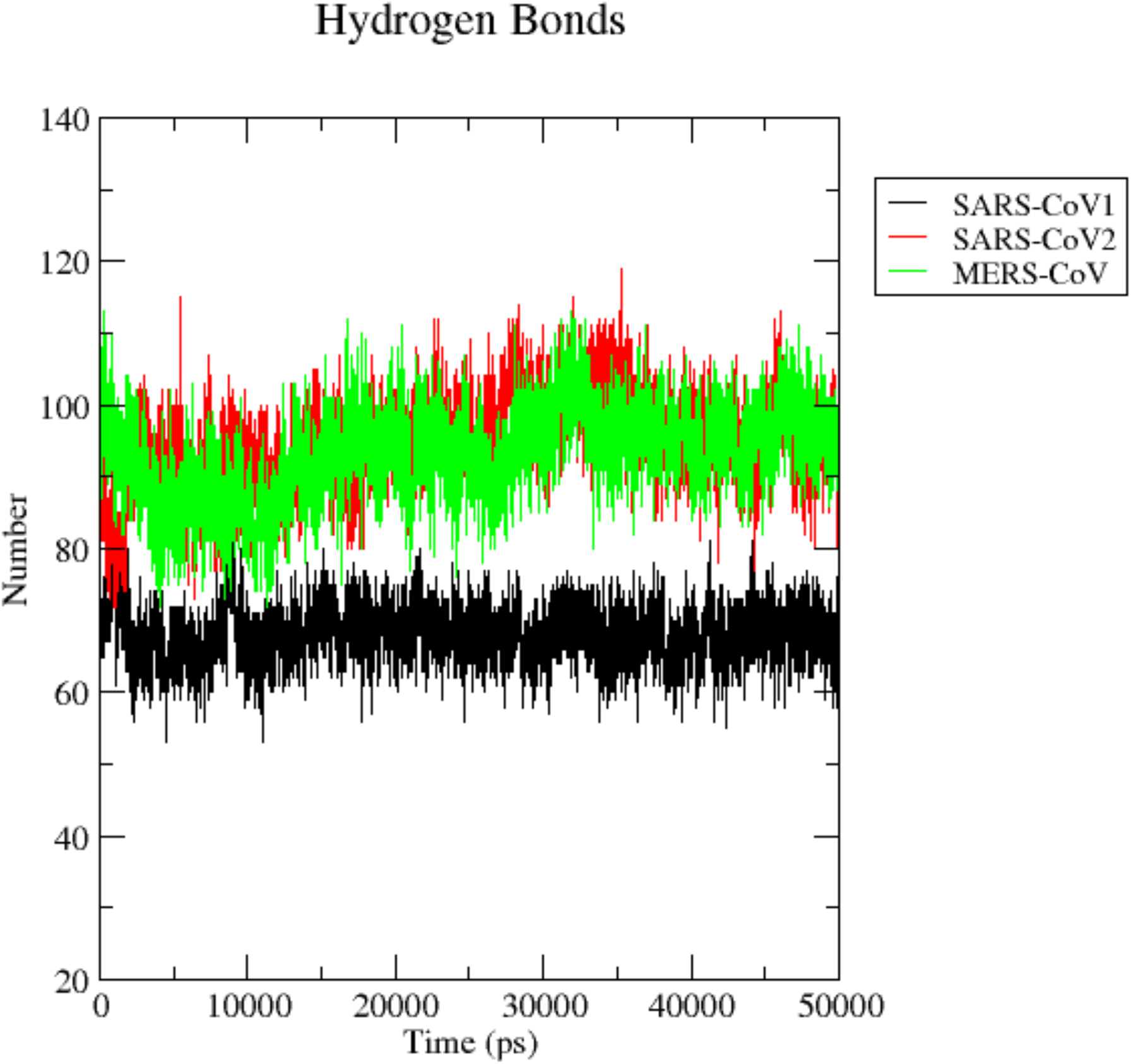
Trajectory analysis of hydrogen bonds. Hydrogen bonds are responsible for the stability of the protein molecules. Black, red and green colour depicts the number of hydrogen bonds of the SARS-CoV1, SARS-CoV2 and MERS-CoV, respectively throughout the simulation run

### 3.4 Ramachandran (ψ/Φ) space of the residues by molecular dynamics Simulation

The frequency of dihedral angles phi (Φ) and psi (Ψ) was monitored during the 50 ns simulation time. The plot of the dihedral angle frequencies in a Ramachandran-like graph provides conformational preferences of the three nsp1 proteins. The allowed regions of the Ramachandran plot of three nsp1 proteins are delimited by green lines in **Figure 5**. High peaks mean that this combination of Ramachandran-angles is assumed often during simulation. In case of SARS-CoV1, the major Ramachandran ψ/Φ angle distribution, as obtained by the MD analysis was found to peak at Ramachandran coordinates of −100° ≤ Φ ≤ −60° and −55° ≤ Ψ ≤ −40°. The other comparatively medium distributions were at −80° ≤ Φ ≤ −60° and 120° ≤ Ψ ≤ 150° in the Ramachandran (ψ, Φ) space. The highly populated Φ/Ψ values were close to the right-handed α-helical space and medium populated Φ/Ψ values were close to the β-sheet space (**Fig.5A and Table 6**). In case of SARS-CoV2, it is worth noting that the distribution of β-sheets (−90° ≤ Φ ≤ −70°, 110° ≤ Ψ ≤ 150°) is more pronounced with respect to the one for α-helices (−95° ≤ Φ ≤ −65°, −40° ≤ Ψ ≤ −30°) (**Fig.5B and Table 6**). The most highly populated (Φ, Ψ) angles for MERS-CoV were −90° ≤ Φ ≤ −65° and 120° ≤ Ψ ≤ 150°, which defined as β-sheet. The other medium and week distributions were at right-handed α-helix and left handed α-helix respectively (**Fig.5C and Table 6**). The analysis of the Ramachandran (ψ/Φ) space of three nsp1 proteins suggests that the distribution of the secondary structures spanned mainly swings between α-helices and β-sheets.

**Table 6:** Ramachandran space of three nsp1 proteins

| Name of the protein | Dihedral angles | Frequency of population | Secondary structure |
| --- | --- | --- | --- |
| 1. SARS-CoV1 | $-100^\circ \leq \Phi \leq -60^\circ$<br>$-55^\circ \leq \Psi \leq -40^\circ$ | Higher | Right-handed α-helix |
| | $-80^\circ \leq \Phi \leq -60^\circ$<br>$120^\circ \leq \Psi \leq 150^\circ$ | Medium | $\beta$ -sheet |
| 2. SARS-CoV2 | $-90^\circ \leq \Phi \leq -70^\circ$<br>$110^\circ \leq \Psi \leq 150^\circ$ | Higher | $\beta$ -sheet |
| | $-95^\circ \leq \Phi \leq -65^\circ$<br>$-40^\circ \leq \Psi \leq -30^\circ$ | Lower | Right-handed α-helix |
| 3. MERS-CoV | $-90^\circ \leq \Phi \leq -65^\circ$<br>$120^\circ \leq \Psi \leq 150^\circ$ | Higher | $\beta$ -sheet |
| | $-90^\circ \leq \Phi \leq -60^\circ$<br>$-40^\circ \leq \Psi \leq -30^\circ$ | Medium | Right-handed α-helix |

**Fig. 5:**
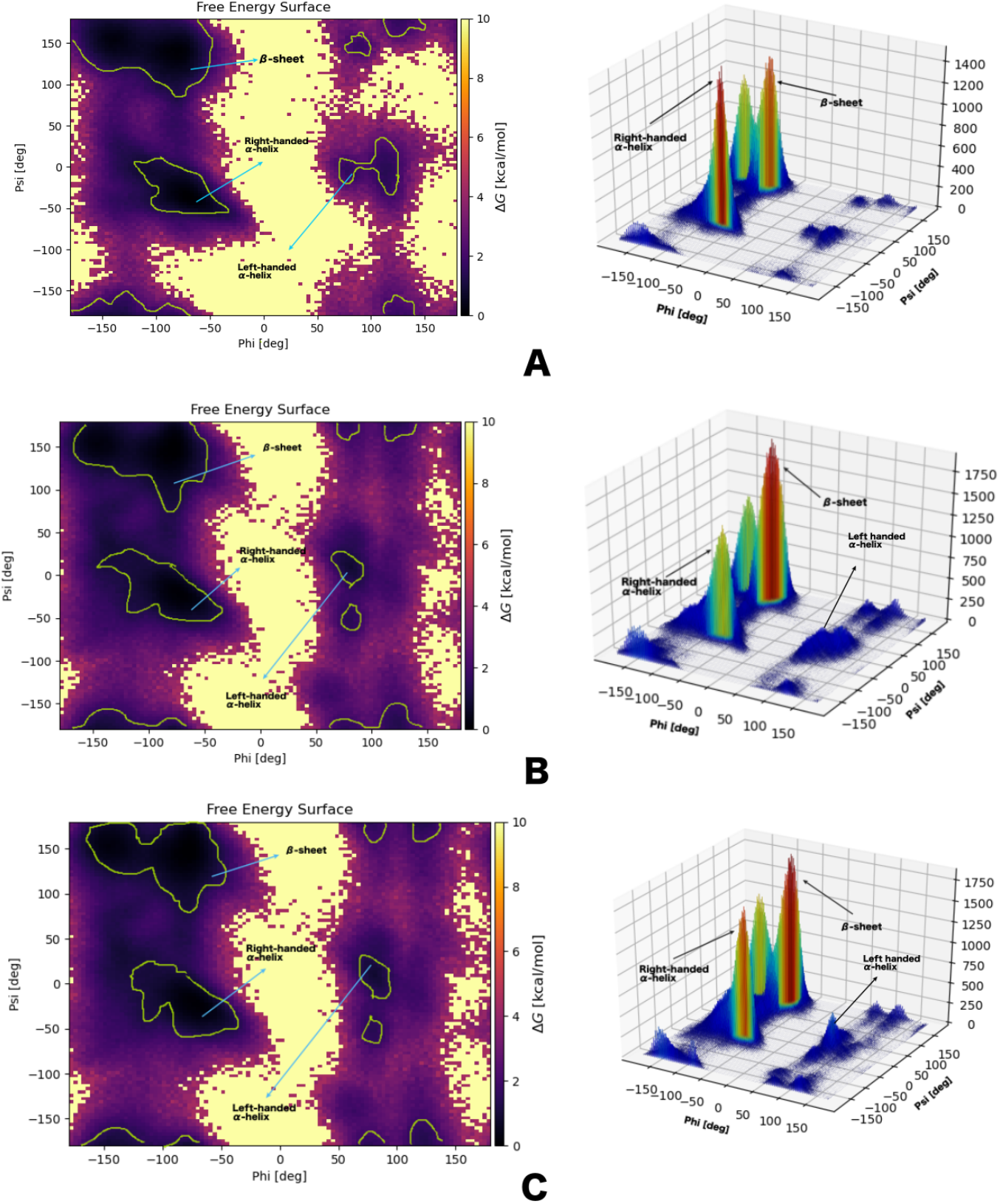
2D and 3D Ramachandran plots obtained by sampling the dihedrals of the three nsp1 during the 50 ns simulations. A. SARS-CoV1. B. SARS-CoV2. C. MERS-CoV. In the 2D plots, the regions delimited by green lines are the most populated and are associated with a well-defined secondary structure. In the 3D plots, the most probable structures are colored in deep orange, while the least probable structures are colored in blue. High peaks mean that this combination of Ramachandran-angles is assumed often during simulation.

### 3.5 Principal component analysis (PCA) and Gibbs free energy landscape (FEL)

The overall pattern of motion of the atoms was monitored using the MD trajectories projected on first (PC1) and second (PC2) principal components to gain a better understanding of the conformational changes in the nsp1 protein of SARS-CoV1, SARS-CoV2 and MERS-CoV. The eigen vectors described the collective motion of the atoms, while the eigenvalues signified the atomic influence in movement. A large distribution of lines indicated greater variance in accordance with more conformational changes in the SARS-CoV2 nsp1 protein compared with SARS-CoV1 and MERS-CoV nsp1 protein. The trajectories of SARS-CoV2 nsp1 protein covered a wider conformational space and showed higher space magnitudes. The trace values, which are the sums of the eigenvalues, were 4.71 nm^2^, 44.082 nm^2^, and 21.611 nm^2^ for SARS-CoV1, SARS-CoV2 and MERS-CoV nsp1, respectively. It was suggested that SARS-CoV2 nsp1 protein appeared to cover a larger conformational space due to its greater flexibility when compared with the other two nsp1 proteins (**Fig. 6**). This observation correlates with the trajectory of the stability parameters such as RMSD, RMSF and Rg of SARS-CoV2 nsp1. It is outward from the PCA plot that the collective motions of the SARS-CoV1 nsp1 protein are localized in a small subspace compared to the SARS-CoV2 and MERS-CoV nsp1 (**Fig.6**)

**Fig. 6:**
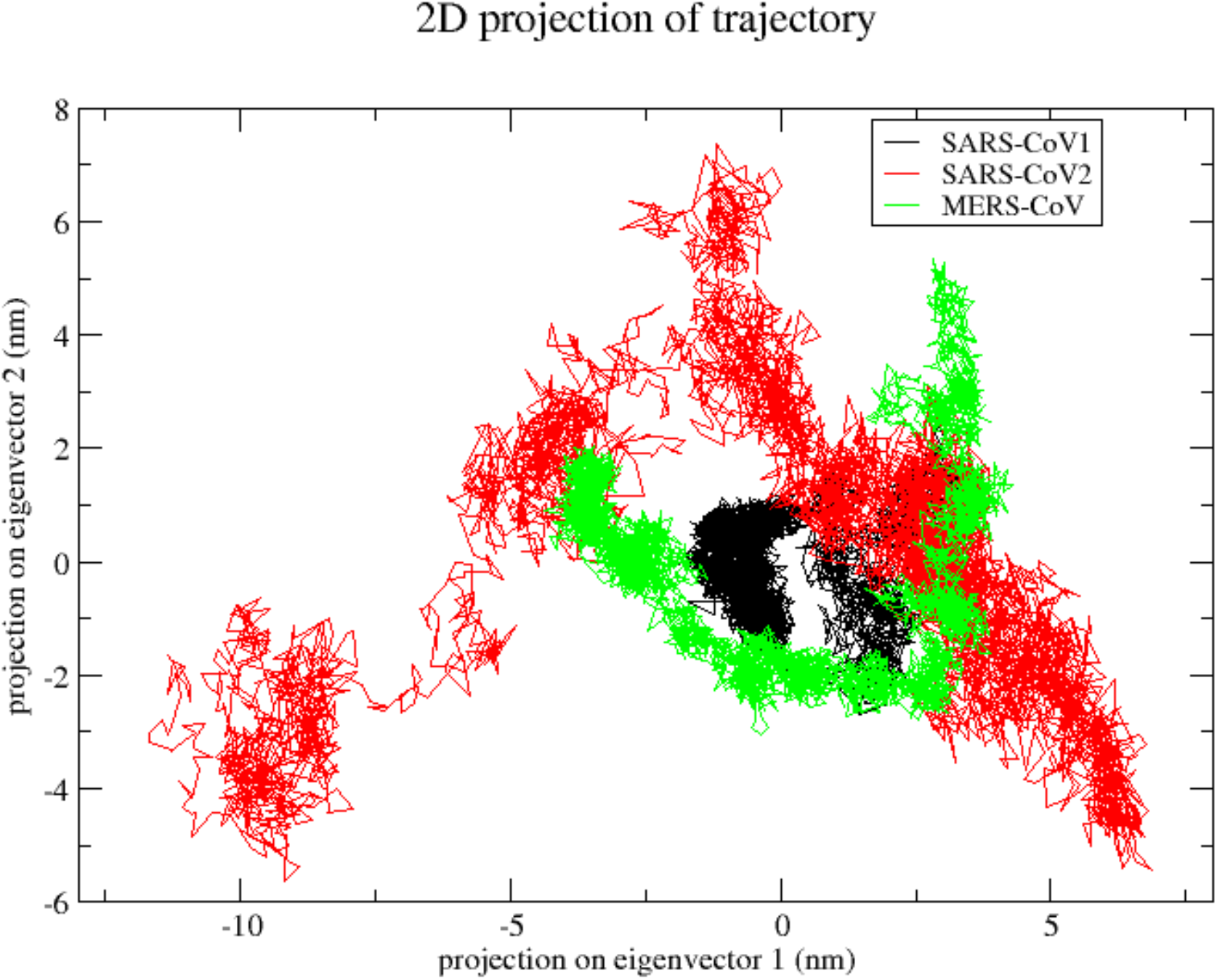
Projection of motion of nsp1 protein atoms of SARS-CoV1 (Black colour), SARS-CoV2 (red colour) and MERS-CoV (green colour) on PC1 and PC2

The Gibbs free energy landscape (FEL) deliver a precise portrayal of a protein’s most stable conformational space, which are certainly important to study the conformational changes during simulation. Gibbs free energy landscape (FEL) was calculated by using PC1, PC2 coordinates and RMSD, Rg coordinates of the three nsp1 protein molecules. Both 2D and 3D graphs of the FEL were plotted using PC1, PC2 and RMSD, Rg coordinates. The ΔG values for SARS-CoV1, SARS-CoV2 and MERS-CoV nsp1 protein were 0 to 13.3, 12.2 and 12.9 kJ/mol respectively. The broader, shallow and narrow energy basin were observed during the trajectory analysis of the simulation. Each nsp1 protein has a different pattern for the FEL. SARS-CoV1 nsp1 has less conformational mobility, restricting to a more confined conformational space within a single local basin with the lowest energy (**Fig.7A**). Extraction of snapshots from this deep wells represents the most native-like conformation which had a RMSD of 0.2702 nm and Rg of 1.2807 nm (**Fig S2A**). In case of SARS-CoV2 nsp1, local minima distributed to about three to four energy basins within the energy landscape which indicates wide range of conformations (**Fig. 7B**). One of the well populated conformations was located at a RMSD of around 0.5433 nm and a Rg of around 1.7332 nm (**Fig.S2B**). MARS-CoV nsp1 has a broader and narrow with two to three energy basins (**Fig. 7C**). The native-like conformation from the low energy basins was centered at the coordinate (RMSD = 0.5724 nm, Rg = 1.6221 nm) (**Fig.S2C**). Figure 7 displays the FELs of SARS-CoV1, SARS-CoV2 and MERS-COV nsp1 proteins where the deeper blue indicates the stable conformational states having lower energy.

**Fig. 7:**
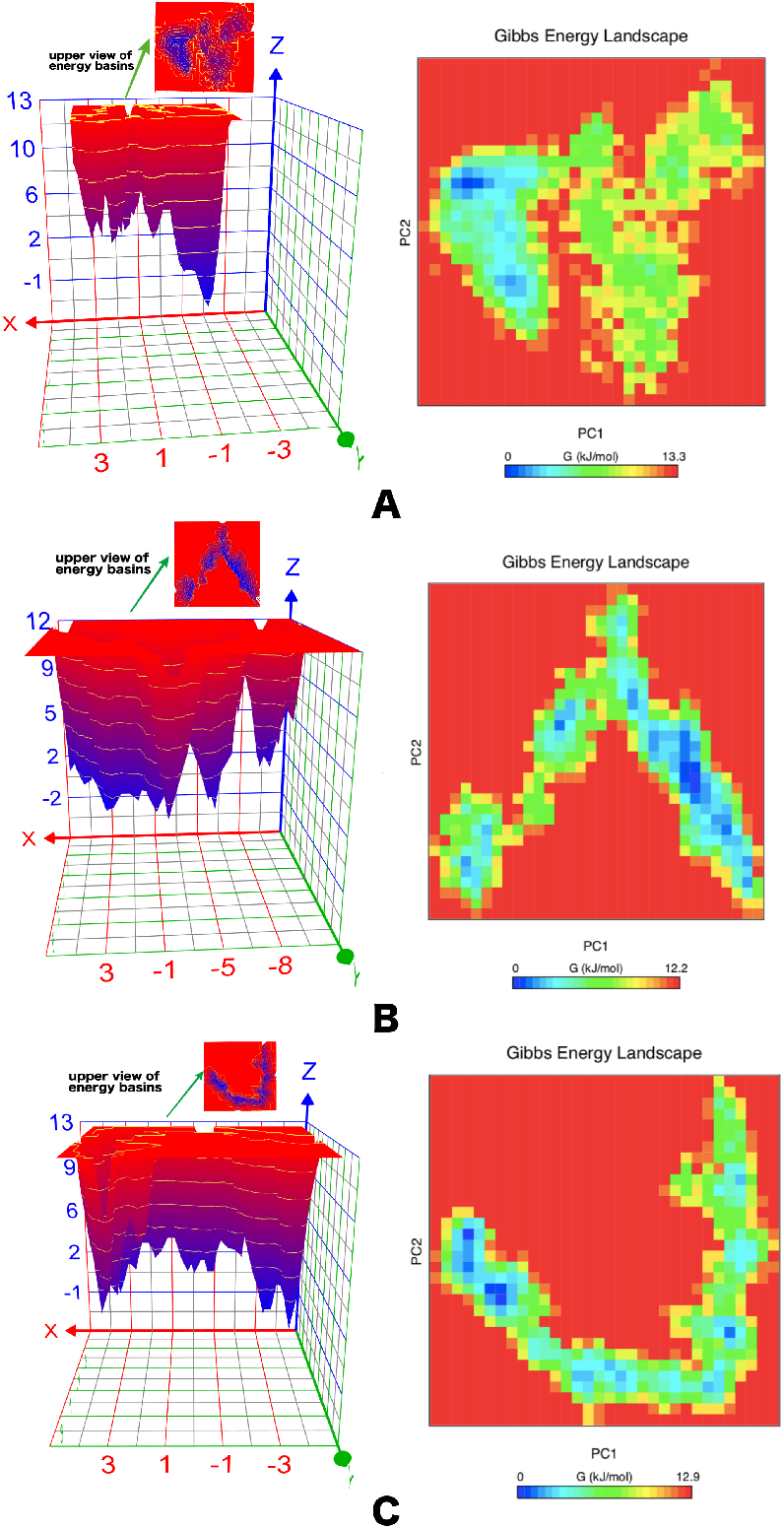
2D and 3D plots of Gibbs free energy landscape (FEL) of three nsp1 proteins. A. SARS-CoV1. B. SARS-CoV2. C. MERS-CoV. The blue, cyan and green regions in the free energy landscape plot denotes low energy state with highly stable protein conformation while the red region signify high energy state with unstable protein conformation

### 3.6 Molecular interactions of SARS-COV2 nsp1 with POLA1

The catalytic domain of DNA polymerase alpha, POLA1 is involved in the replication process. The molecular association of SARS-COV2 nsp1 with POLA1 was predicted by the HADDOCK programme. The nsp1-POLA1 docked complexes were analyzed based on Z-score and HAD-DOCK score (**Fig. 8 & Table S2**). PRODIGY was used to predict the binding energy for each nsp1-POLA1 complex from the best there cluster. Three best docked complexes were selected on the basis of lowest binding energy. (**Fig. S3**). The best energy values obtained were −13.0 kcal/ mol, −11.6 kcal/mol and −9.6 kcal/mol. **(Table S2)**. Interface area, involvement of amino acids and molecular interactions were calculated by PISA server. The interface area of the best docked complex (ΔG = −13.0 kcal/mol) was 1260.9 Å^2^ and shown in **Fig. 9A**. The best complex structure of SARS-COV2 nsp1-POLA1 was evaluated by the Ramachandran plot which shows 98.4% residues are present in the allowed region. From ProSA and ProQ analysis, it is clear that the overall model quality of the final protein–protein complex is within the range of scores typically found for proteins of similar size (**Table S1**). Interaction studies of this nsp1-POLA1 complex showed thirteen hydrogen bonds and eight salt bridge interactions at the interface region (**Table 7** & **Fig. 9B**). It was observed from the docking experiments that the residues of finger domain (Lys923, Gln927, Gln932) and palm domain (Glu1060, Lys1020, Lys1024, Asn1030, Lys1031 and Glu1037) of POLA1 are mainly involved in the binding process with SARS-COV2 nsp1 protein (**Fig. 8**). The hydrogen bonds and salt bridges interactions play an important role towards the stability of the SARS-COV2 nsp1-POLA1 complex formation.

**Table 7:** Intermolecular interactions of the best docked complex of SARS-CoV2 nsp1-POLA1 predicted by PISA analysis

| S l. No | Amino acids of nsp1 | Amino acids of POLA1 | Distance (Å) | Interactions |
| --- | --- | --- | --- | --- |
| 1 | Glu 2 [OE2] | Lys 923 [HZ2] | 1.63 | Hydrogen bond |
| 2 | Lys47 [O] | Lys 923 [HZ1] | 1.82 | Hydrogen bond |
| 3 | Glu 2 [OE1] | Gln 927 [HE21] | 1.81 | Hydrogen bond |
| 4 | Met 1 [O] | Gln 927 [HE22] | 2.17 | Hydrogen bond |
| 5 | Pro 115 [O] | Gln 932 [HE21] | 1.78 | Hydrogen bond |
| 6 | Asp 33 [OD2] | Lys 1020 [HZ1] | 1.62 | Hydrogen bond |
| 7 | Glu 41 [OE1] | Ans 1030 [HD21] | 1.78 | Hydrogen bond |
| 8 | Glu 36 [O] | Lys 1031 [HZ3] | 1.80 | Hydrogen bond |
| 9 | Met 1 [N] | Gln 927 [OE1] | 3.50 | Hydrogen bond |
| 10 | Arg 29 [HH21] | Glu 1016 [OE2] | 2.11 | Hydrogen bond |
| 11 | Arg 29 [HH11] | Glu 1016 [OE2] | 1.94 | Hydrogen bond |
| 12 | Lys47 [HZ3] | Glu 1037 [OE1] | 1.63 | Hydrogen bond |
| 13 | Lys47 [HZ1] | Glu 1037 [OE2] | 2.36 | Hydrogen bond |
| 14 | Glu 2 [OE2] | Lys 923 [NZ] | 2.57 | Salt bridge |
| 15 | Asp 33 [OD2] | Lys 1020 [NZ] | 2.64 | Salt bridge |
| 16 | Glu 37 [OE1] | Lys 1024 [NZ] | 2.88 | Salt bridge |
| 17 | Glu 37 [OE2] | Lys 1024 [NZ] | 2.91 | Salt bridge |
| 18 | Arg 29 [NH2] | Glu 1016 [OE2] | 2.97 | Salt bridge |
| 19 | Arg 29 [NH1] | Glu 1016 [OE2] | 2.85 | Salt bridge |
| 20 | Lys47 [NZ] | Glu 1037 [OE1] | 2.67 | Salt bridge |
| 21 | Lys47 [NZ] | Glu 1037 [OE2] | 2.70 | Salt bridge |

**Fig. 8:**
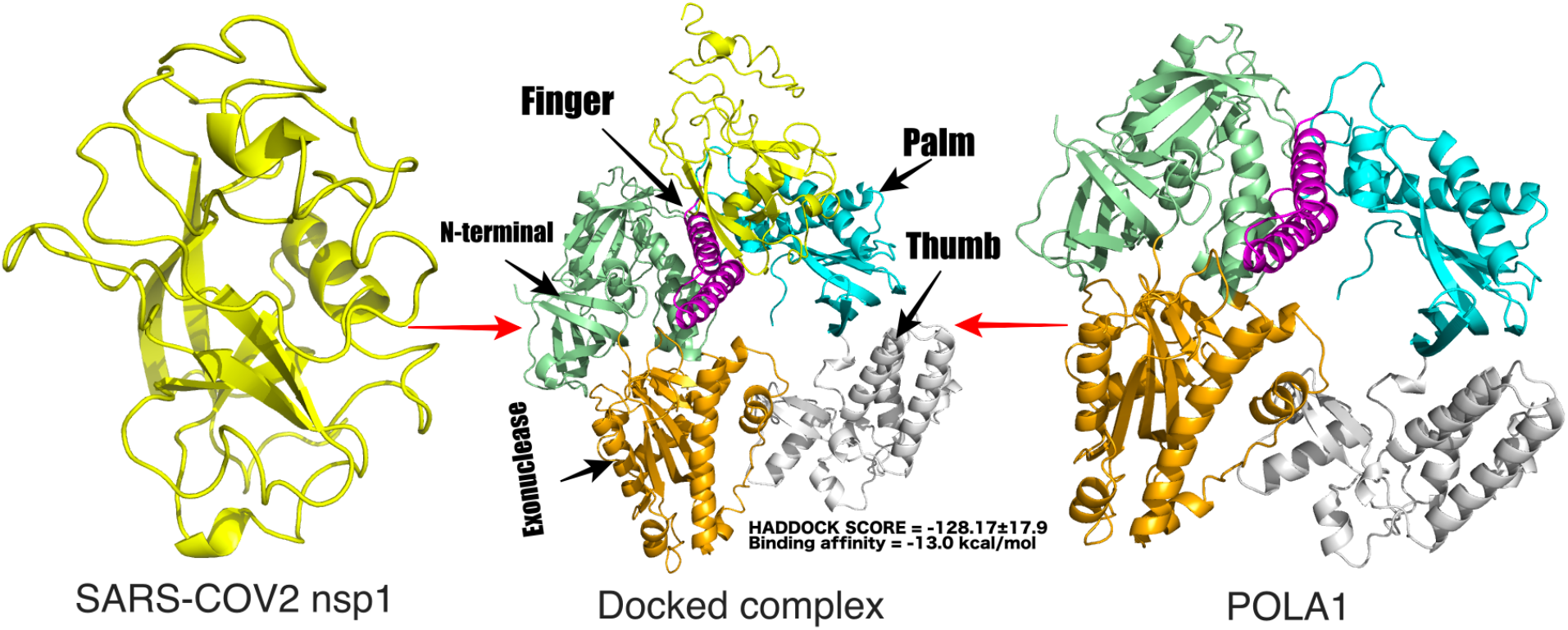
The structure of docking complex between SARS-CoV2 nsp1 protein and POLA1. SARS-CoV2 nsp1 is represented by a yellow cartoon. POLA1 is composed of five domains. N-terminal (338-534 & 761-808), exonuclease (535-760), palm (834-908 & 968-1076), finger (909-967), and thumb (1077-1250) domain are represented by pale green, orange, cyan, magenta and grey colour respectively. The binding affinity of nsp1 is higher at the interface region of palm (cyan colour) and finger (magenta colour) domain of catalytic subunit of DNA polymerase alpha POLA1

**Fig. 9:**
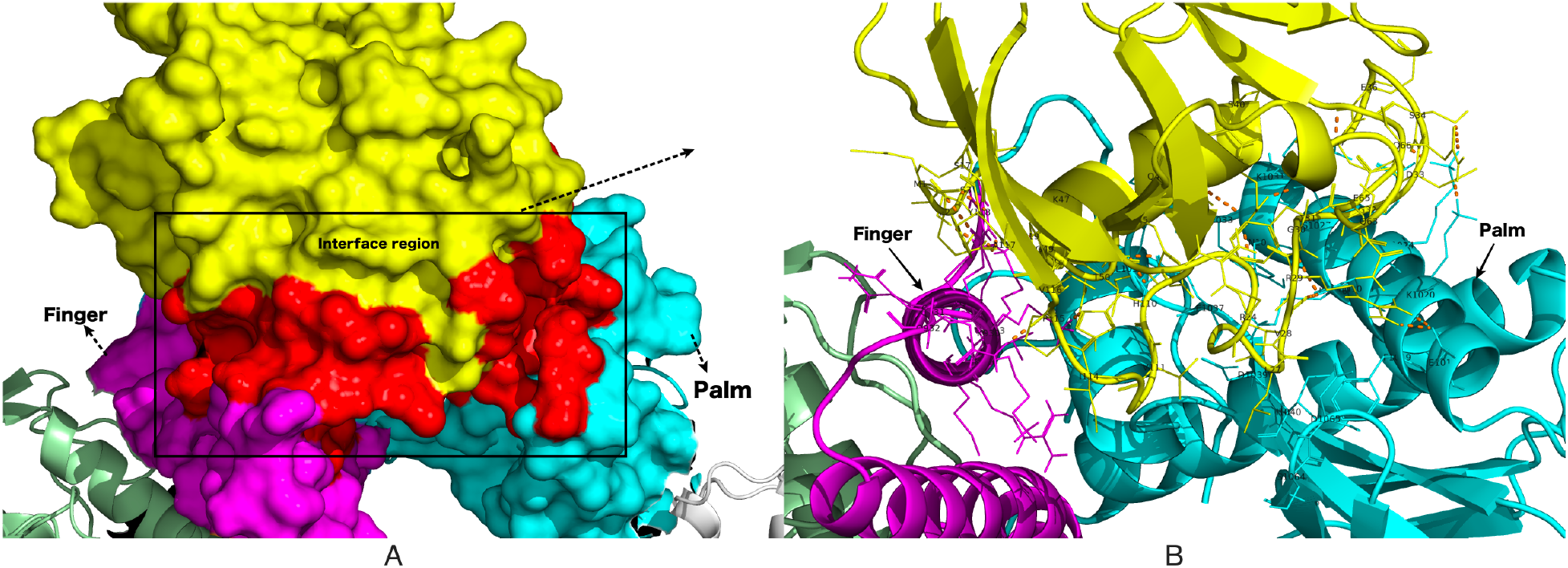
The proposed binding mode of the host cell POLA1 and the COVID-19 nsp1 model. A. Nsp1 (yellow surface) interacts with the palm (cyan surface) and finger (magenta surface) domain. Interface region is represented by a red surface. B. Molecular interactions between SARS-CoV2 nsp1 and POLA1. Interface residues are represented as a line model. Several bonds are depicted by orange dotted line

## 4. Conclusion

COVID-19 pandemic leads to a health, economic, and social crisis in the world. The development of a specific targeted therapy could reduce the rate of infection. This comprehensive study represents an immunoinformatics approach towards the identification of specific B-cell and T-cell epitopes of three nsp1 proteins. Four epitope rich regions (15-27, 45-81, 121-140 and 147-178) that were shared between T-cell and B-cell were reported in SARS-CoV2 nsp1 protein. The in-depth structural elucidation of nsp1 proteins together with dynamic conformations showed that SARS-CoV2 nsp1 protein covers a large conformational space due to its greater flexibility compared with SARS-CoV1 and MERS-CoV nsp1. A three-dimensional structural model of the complex structure between SARS-CoV2 nsp1 protein and catalytic subunit of DNA polymerase alpha POLA1 was constructed using protein-protein docking approach. During complex formation between SARS-CoV2 nsp1 and POLA1, salt bridge interactions help to bring the two proteins in close proximity and form 13 strong hydrogen bonds that contribute to the stability of the complex formation. Knowledge of this important binding site could open the door for further simulation and experimental studies on the mode of SARS-CoV2 nsp1 protein recognition by the catalytic site of DNA polymerase alpha POLA1. From FEL analysis, it was observed that as SARS-CoV1 and MERS-CoV nsp1 has shown stable RMSD and Rg so their energy minimas are confined in a particular region. Taken all together, according to structural evaluation as well as immunological analysis, nsp1 protein could be considered as a possible drug target and candidate molecule for the vaccine development process against COVID-19.

## Acknowledgement

The author wishes to express sincere thanks to Dr. Sibani Sen Chakraborty, West Bengal State University, Kolkata and Dr. Asim Kumar Bera, Institute for Protein Design, Seattle, for their useful suggestions in preparation of the manuscript.

## Data Availability

The modelled and docking structures are available upon request from the corresponding author.

## CRediT authorship contribution statement

**Ankur Chaudhuri**: Supervision, Conceptualization, Formal analysis, Writing - review & editing.

## Highlights

- Structural elucidation at molecular level of nsp1 of SARS-CoV1, SARS-CoV2, and MERS-CoV
- Identifications of epitopes by immunoinformatics approach
- SARS-CoV2 nsp1 cover a large conformational space due to greater flexibility
- Molecular docking between SARS-CoV2 nsp1 and POLA1 to identify important residues
- Structural insights of nsp1 could be used in drug design process against COVID-19

## Graphical abstract

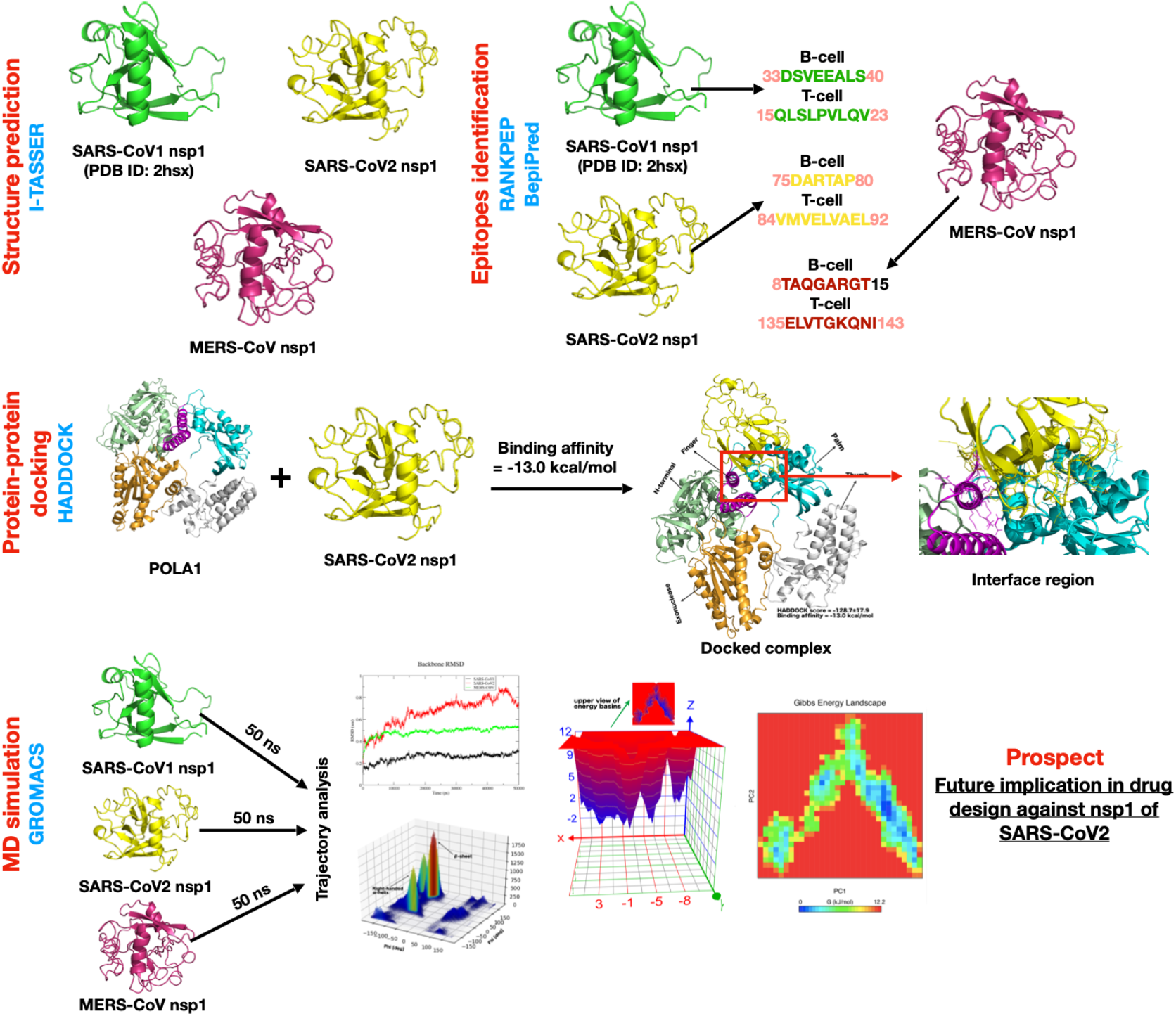

## Supplementary file

**Fig. S1:**
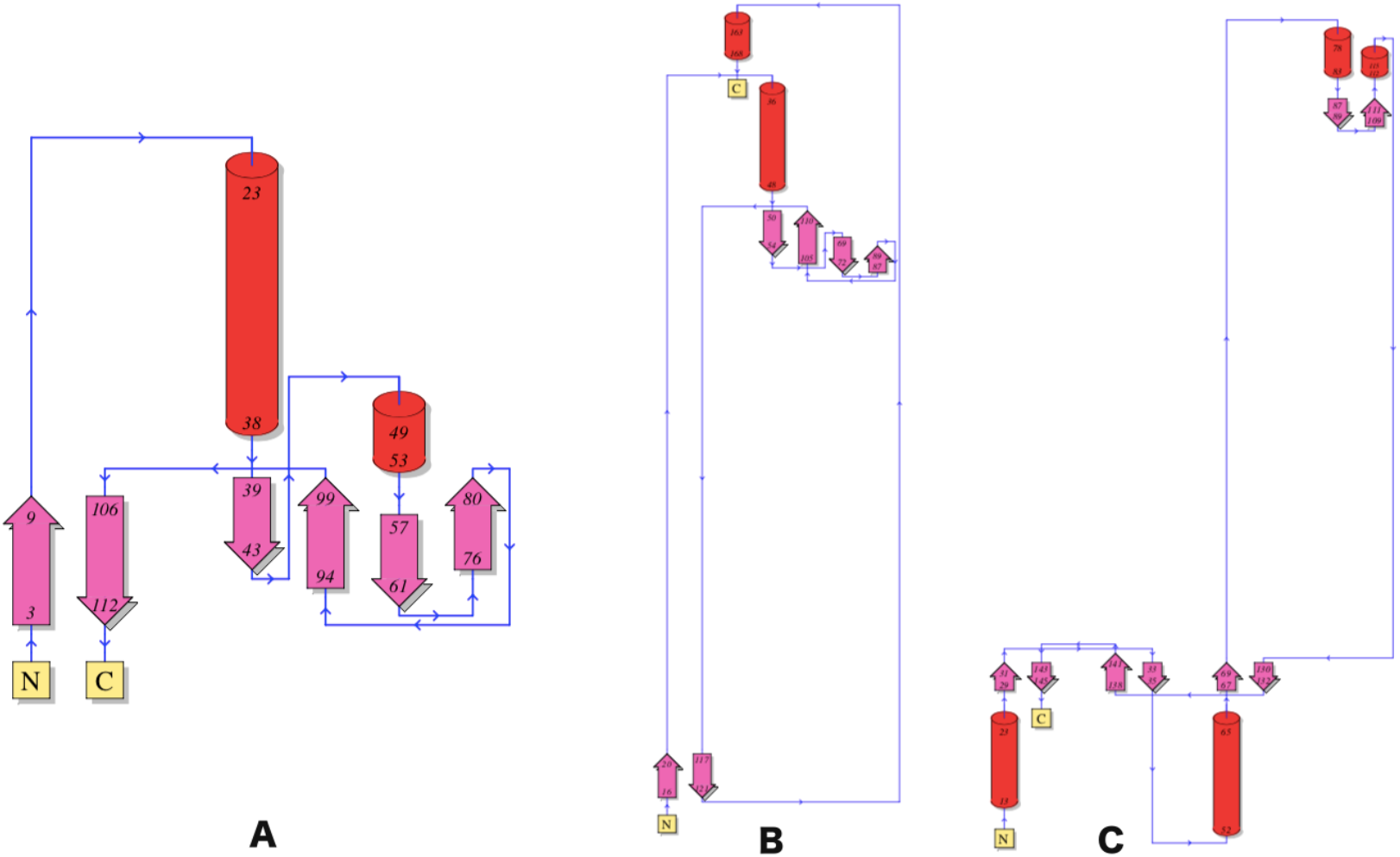
Topological analysis by using PDBsum database. A. SARS-CoV1 nsp1. B. SARS-CoV2 nsp1. C. MERS-CoV nsp1

**Fig. S2:**
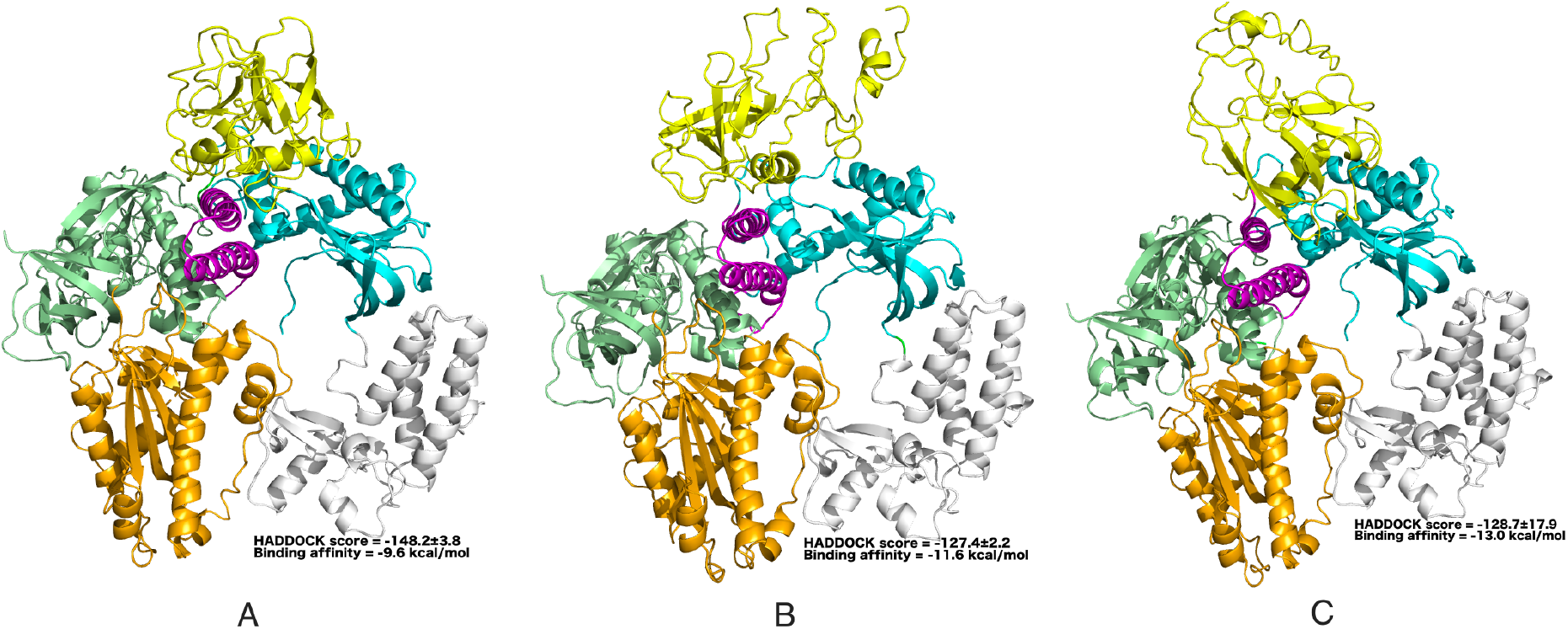
Best SARS-CoV2 nsp1-POLA1 complex model. A. Top model from Cluster 1. B. Top model from cluster 2. C. Top model from cluster 5. Best model from each cluster was predicted according to lowest binding energy. Binding affinity is calculated by PRODIGY server

**Fig. S3:**
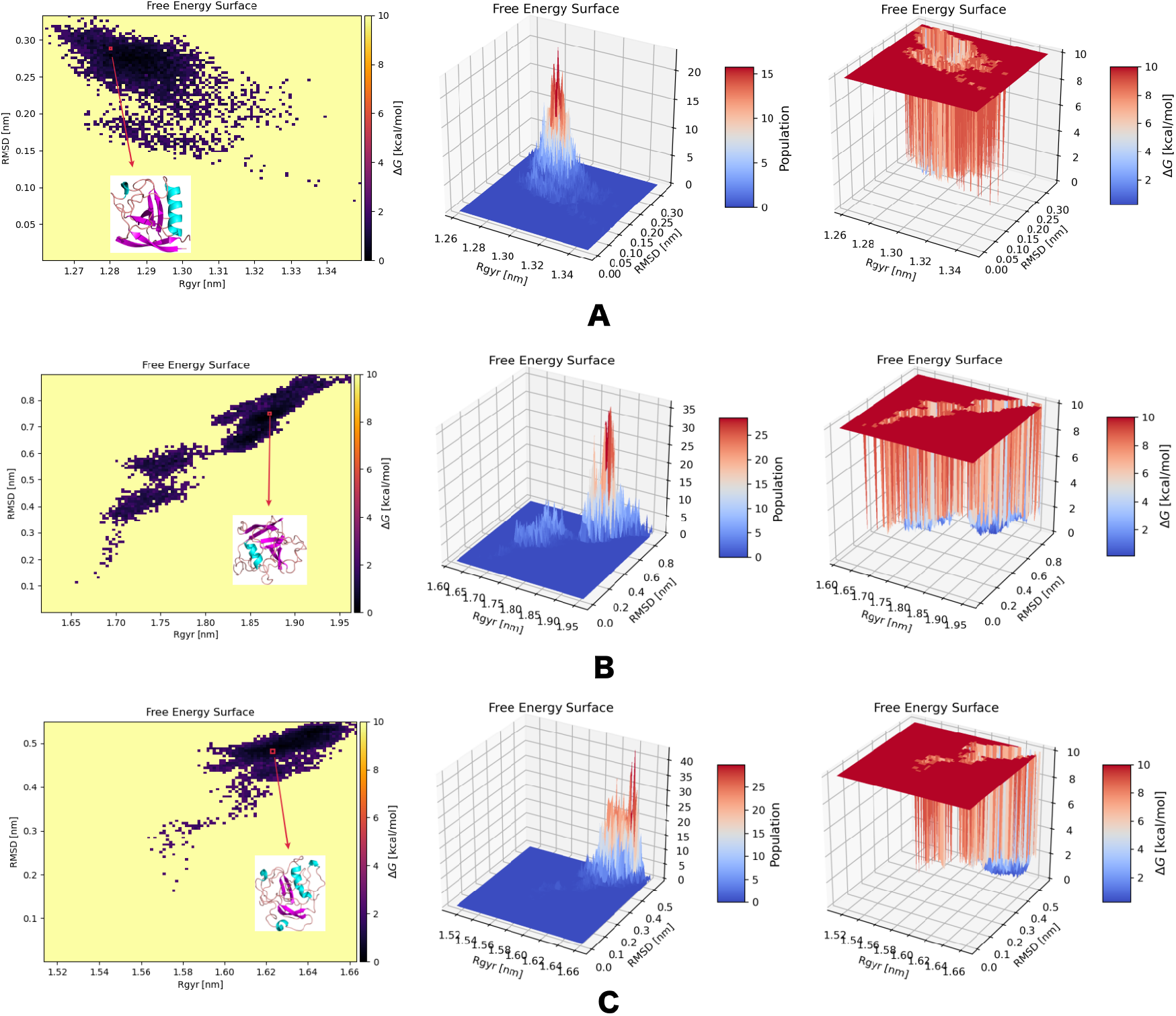
2D and 3D plots of Gibbs free energy landscape (FEL) of three nsp1 proteins by using rmsd and Rg coordinates **A. SARS-CoV1. B. SARS-CoV2. C. MERS-CoV**.

**Table S1:** Three-dimensional model validation using several analyses

| **Model Validation** | **SARS-CoV2 nsp1** | **MERS-CoV nsp1** | **SARS-CoV2 nsp1-POLA1 complex** |
| --- | --- | --- | --- |
| Ramachandran plot | 93.95% residues in allowed region | 92.07% residues in allowed region | 98.4% residues in allowed region |
| ProSA | Z-score = -4.78 | Z-score = -2.42 | Z-score = -13.55 |
| ProQ* | LG score = 3.578 | LG score = 3.607 | LG score = 4.418 |

\*Notes: LGscore > 1.5 fairly good model, LGscore > 2.5 very good model, LGscore > 4 extremely good model.

**Table S2:**
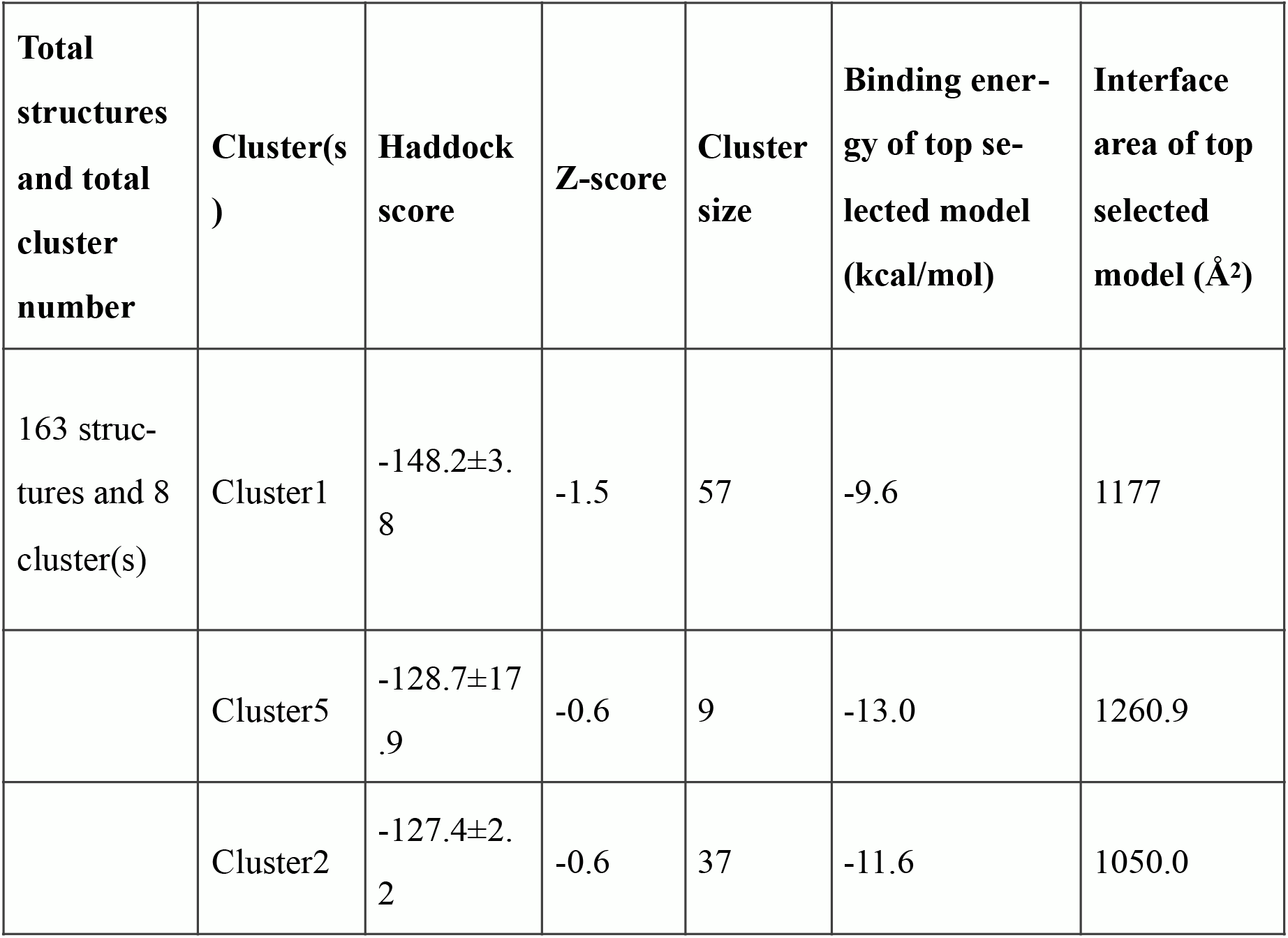
Protein-protein docking between SARS-CoV2 nsp1 and POLA1 by HADDOCK server

| **Total structures and total cluster number** | **Cluster(s)** | **Haddock score** | **Z-score** | **Cluster size** | **Binding energy of top selected model (kcal/mol)** | **Interface area of top selected model (Å<sup>2</sup>)** |
| --- | --- | --- | --- | --- | --- | --- |
| 163 structures and 8 cluster(s) | Cluster1 | -148.2±3.8 | -1.5 | 57 | -9.6 | 1177 |
|  | Cluster5 | -128.7±17.9 | -0.6 | 9 | -13.0 | 1260.9 |
|  | Cluster2 | -127.4±2.2 | -0.6 | 37 | -11.6 | 1050.0 |

## References

1. R. Lu, X. Zhao, J. Li, P. Niu, B. Yang, H. Wu, et al., Genomic characterisation and epidemiology of 2019 novel coronavirus: implications for virus origins and receptor binding, The Lancet. 395 (2020) 565–574. doi:10.1016/s0140-6736(20)30251-8

2. F. Wu, S. Zhao, B. Yu, Y.-M. Chen, W. Wang, Z.-G. Song, et al., A new coronavirus associated with human respiratory disease in China, Nature. 579 (2020) 265–269. doi:10.1038/s41586-020-2008-3

3. (worldometers.info. (2020). COVID-19 coronavirus pandemic. https://www.worldometers.in-fo/coronavirus/ (Last accessed May 25, 2020)

4. A.A. Elfiky, S.M. Mahdy, W.M. Elshemey, Quantitative structure-activity relationship and molecular docking revealed a potency of anti-hepatitis C virus drugs against human corona viruses, Journal of Medical Virology. 89 (2017) 1040–1047. doi:10.1002/jmv.24736

5. I.I. Bogoch, A. Watts, A. Thomas-Bachli, C. Huber, M.U.G. Kraemer, K. Khan, Potential for global spread of a novel coronavirus from China, Journal of Travel Medicine. 27 (2020). doi: 10.1093/jtm/taaa011

6. P.C.Y. Woo, Y. Huang, S.K.P. Lau, K.-Y. Yuen, Coronavirus Genomics and Bioinformatics Analysis, Viruses. 2 (2010) 1804–1820. doi:10.3390/v2081803

7. S.F. Ahmed, A.A. Quadeer, M.R. Mckay, Preliminary Identification of Potential Vaccine Targets for the COVID-19 Coronavirus (SARS-CoV-2) Based on SARS-CoV Immunological Studies, Viruses. 12 (2020) 254. doi:10.3390/v12030254

8. D.E. Gordon, G.M. Jang, M. Bouhaddou, J. Xu, K. Obernier, K.M. White, et al., A SARS-CoV-2 protein interaction map reveals targets for drug repurposing, Nature. 583 (2020) 459–468. doi:10.1038/s41586-020-2286-9

9. B.W. Neuman, P. Chamberlain, F. Bowden, J. Joseph, Atlas of coronavirus replicase structure, Virus Research. 194 (2014) 49–66. doi:10.1016/j.virusres.2013.12.004

10. R.F. Connor, R.L. Roper, Unique SARS-CoV protein nsp1: bioinformatics, biochemistry and potential effects on virulence, Trends in Microbiology. 15 (2007) 51–53. doi:10.1016/j.tim.2006.12.005

11. W. Kamitani, C. Huang, K. Narayanan, K.G. Lokugamage, S. Makino, A two-pronged strate-gy to suppress host protein synthesis by SARS coronavirus Nsp1 protein, Nature Structural &amp; Molecular Biology. 16 (2009) 1134–1140. doi:10.1038/nsmb.1680

12. K.G. Lokugamage, K. Narayanan, C. Huang, S. Makino, Severe Acute Respiratory Syndrome Coronavirus Protein nsp1 Is a Novel Eukaryotic Translation Inhibitor That Represses Multiple Steps of Translation Initiation, Journal of Virology. 86 (2012) 13598–13608. doi:10.1128/jvi.01958-12

13. R. Channappanavar, A.R. Fehr, R. Vijay, M. Mack, J. Zhao, D.K. Meyerholz, et al., Dysregulated Type I Interferon and Inflammatory Monocyte-Macrophage Responses Cause Lethal Pneumonia in SARS-CoV-Infected Mice, Cell Host &amp; Microbe. 19 (2016) 181–193. doi: 10.1016/j.chom.2016.01.007

14. E. Kindler, V. Thiel, SARS-CoV and IFN: Too Little, Too Late, Cell Host & Microbe. 19 (2016) 139–141. doi:10.1016/j.chom.2016.01.012

15. A.H.Y. Law, D.C.W. Lee, B.K.W. Cheung, H.C.H. Yim, A.S.Y. Lau, Role for Nonstructural Protein 1 of Severe Acute Respiratory Syndrome Coronavirus in Chemokine Dysregulation, Journal of Virology. 81 (2006) 416–422. doi:10.1128/jvi.02336-05

16. NCBI. National Center of Biotechnology Informatics (NCBI) database website http://www.ncbi.nlm.nih.gov/. 2020. http://www.ncbi.nlm.nih.gov/2020 (xLast accessed May 25, 2020)

17. W. Gish, D.J. States, Identification of protein coding regions by database similarity search, Nature Genetics. 3 (1993) 266–272. doi:10.1038/ng0393-266

18. C. Notredame, D.G. Higgins, J. Heringa, T-coffee: a novel method for fast and accurate multiple sequence alignment 1 1 Edited by J. Thornton, Journal of Molecular Biology. 302 (2000) 205–217. doi:10.1006/jmbi.2000.4042

19. M.S. Almeida, M.A. Johnson, T. Herrmann, M. Geralt, Wüthrich Kurt, Novel β-Barrel Fold in the Nuclear Magnetic Resonance Structure of the Replicase Nonstructural Protein 1 from the Severe Acute Respiratory Syndrome Coronavirus, Journal of Virology. 81 (2007) 3151–3161. doi:10.1128/jvi.01939-06

20. J. Yang, R. Yan, A. Roy, D. Xu, J. Poisson, Y. Zhang, The I-TASSER Suite: protein structure and function prediction, Nature Methods. 12 (2014) 7–8. doi:10.1038/nmeth.3213

21. W. Zheng, C. Zhang, Q. Wuyun, R. Pearce, Y. Li, Y. Zhang, LOMETS2: improved meta-threading server for fold-recognition and structure-based function annotation for distant-homology proteins, Nucleic Acids Research. 47 (2019). doi:10.1093/nar/gkz384

22. D. Cozzetto, A. Kryshtafovych, A. Tramontano, Evaluation of CASP8 model quality predictions, Proteins: Structure, Function, and Bioinformatics. 77 (2009) 157–166. doi:10.1002/prot.22534

23. R.A. Laskowski, M.W. Macarthur, D.S. Moss, J.M. Thornton, PROCHECK: a program to check the stereochemical quality of protein structures, Journal of Applied Crystallography. 26 (1993) 283–291. doi:10.1107/s0021889892009944

24. A.L. Morris, M.W. Macarthur, E.G. Hutchinson, J.M. Thornton, Stereochemical quality of protein structure coordinates, Proteins: Structure, Function, and Genetics. 12 (1992) 345–364. doi:10.1002/prot.340120407

25. M. Wiederstein, M.J. Sippl, ProSA-web: interactive web service for the recognition of errors in three-dimensional structures of proteins, Nucleic Acids Research. 35 (2007). doi:10.1093/nar/gkm290

26. S. Cristobal, A. Zemla, D. Fischer, L. Rychlewski, A. Elofsson, A study of quality measures for protein threading models, BMC Bioinformatics. 2 (2001) 5. doi:10.1186/1471-2105-2-5

27. J. Yang, A. Roy, Y. Zhang, Protein–ligand binding site recognition using complementary bind-ing-specific substructure comparison and sequence profile alignment, Bioinformatics. 29 (2013) 2588–2595. doi:10.1093/bioinformatics/btt447

28. J. Yang, A. Roy, Y. Zhang, BioLiP: a semi-manually curated database for biologically relevant ligand–protein interactions, Nucleic Acids Research. 41 (2012). doi:10.1093/nar/gks966

29. P.A. Reche, E.L. Reinherz, Prediction of Peptide-MHC Binding Using Profiles, Methods in Molecular Biology Immunoinformatics. (2007) 185–200. doi:10.1007/978-1-60327-118-9_13

30. C. Kruiswijk, G. Richard, M.L. Salverda, P. Hindocha, W.D. Martin, A.S.D. Groot, et al., In silico identification and modification of T cell epitopes in pertussis antigens associated with tolerance, Human Vaccines &amp; Immunotherapeutics. 16 (2020) 277–285. doi: 10.1080/21645515.2019.1703453

31. R. Vita, S. Mahajan, J.A. Overton, S.K. Dhanda, S. Martini, J.R. Cantrell, et al., The Immune Epitope Database (IEDB): 2018 update, Nucleic Acids Research. 47 (2018). doi:10.1093/nar/gky1006

32. A. Kolaskar, P.C. Tongaonkar, A semi-empirical method for prediction of antigenic determi-nantsonproteinantigens, FEBS Letters. 276 (1990) 172–174. doi:10.1016/0014-5793(90)80535-q

33. M.U. Mirza, S. Rafique, A. Ali, M. Munir, N. Ikram, A. Manan, et al., Towards peptide vaccines against Zika virus: Immunoinformatics combined with molecular dynamics simulations to predict antigenic epitopes of Zika viral proteins, Scientific Reports. 6 (2016). doi:10.1038/srep37313

34. H. Berendsen, D.V.D. Spoel, R.V. Drunen, GROMACS: A message-passing parallel molecular dynamics implementation, Computer Physics Communications. 91 (1995) 43–56. doi: 10.1016/0010-4655(95)00042-e

35. W.L. Jorgensen, D.S. Maxwell, J. Tirado-Rives, Development and Testing of the OPLS All-Atom Force Field on Conformational Energetics and Properties of Organic Liquids, Journal of the American Chemical Society. 118 (1996) 11225–11236. doi:10.1021/ja9621760

36. W.L. Jorgensen, J. Chandrasekhar, J.D. Madura, R.W. Impey, M.L. Klein, Comparison of simple potential functions for simulating liquid water, The Journal of Chemical Physics. 79 (1983) 926–935. doi:10.1063/1.445869

37. M. Parrinello, A. Rahman, Polymorphic transitions in single crystals: A new molecular dynamics method, Journal of Applied Physics. 52 (1981) 7182–7190. doi:10.1063/1.328693

38. U. Essmann, L. Perera, M.L. Berkowitz, T. Darden, H. Lee, L.G. Pedersen, A smooth particle mesh Ewald method, The Journal of Chemical Physics. 103 (1995) 8577–8593. doi: 10.1063/1.470117

39. B. Hess, H. Bekker, H.J.C. Berendsen, J.G.E.M. Fraaije, LINCS: A linear constraint solver for molecular simulations, Journal of Computational Chemistry. 18 (1997) 1463–1472. doi: 10.1002/(sici)1096-987x(199709)18:12<1463::aid-jcc4>3.0.co;2-h

40. W. Humphrey, A. Dalke, K. Schulten, VMD: Visual molecular dynamics, Journal of Molecular Graphics. 14 (1996) 33–38. doi:10.1016/0263-7855(96)00018-5

41. E.F. Pettersen, T.D. Goddard, C.C. Huang, G.S. Couch, D.M. Greenblatt, E.C. Meng, et al., UCSF Chimera? A visualization system for exploratory research and analysis, Journal of Computational Chemistry. 25 (2004) 1605–1612. doi:10.1002/jcc.20084

42. WL DeLano, PyMOL: an open-source molecular graphics tool. Ccp4 Newslett Protein Crystallogr 2002;40:11

43. PJ Turner, XMGRACE, Version 5.1.19. 2005; Central for costal and Land-Margin Research; Oregon Graduate Institute of Science and Technology, Beaverton, ORE, USA

44. A. Amadei, A.B.M. Linssen, H.J.C. Berendsen, Essential dynamics of proteins, Proteins: Structure, Function, and Genetics. 17 (1993) 412–425. doi:10.1002/prot.340170408

45. H. Frauenfelder, S. Sligar, P. Wolynes, The energy landscapes and motions of proteins, Science. 254 (1991) 1598–1603. doi:10.1126/science.1749933

46. G.V. Zundert, J. Rodrigues, M. Trellet, C. Schmitz, P. Kastritis, E. Karaca, et al., The HAD-DOCK2.2 Web Server: User-Friendly Integrative Modeling of Biomolecular Complexes, Journal of Molecular Biology. 428 (2016) 720–725. doi:10.1016/j.jmb.2015.09.014

47. T. Tahirov, A. Baranovskiy, N. Babayeva, Crystal Structure Of The Catalytic Core Of Human Dna Polymerase Alpha In Ternary Complex With An Dna-Primed Dna Template And Dctp, (2018). doi:10.2210/pdb6as7/pdb

48. L.C. Xue, J.P. Rodrigues, P.L. Kastritis, A.M. Bonvin, A. Vangone, PRODIGY: a web server for predicting the binding affinity of protein–protein complexes, Bioinformatics. (2016). doi: 10.1093/bioinformatics/btw514

49. M. Thoms, R. Buschauer, M. Ameismeier, L. Koepke, T. Denk, M. Hirschenberger, et al., Structural basis for translational shutdown and immune evasion by the Nsp1 protein of SARS-CoV-2, Science. 369 (2020) 1249–1255. doi:10.1126/science.abc8665

50. R. Channappanavar, J. Zhao, S. Perlman, T cell-mediated immune response to respiratory coronaviruses, Immunologic Research. 59 (2014) 118–128. doi:10.1007/s12026-014-8534-z

51. H.-L.J. Oh, S.K.-E. Gan, A. Bertoletti, Y.-J. Tan, Understanding the T cell immune response in SARS coronavirus infection, Emerging Microbes &amp; Infections. 1 (2012) 1–6. doi: 10.1038/emi.2012.26

52. P.-R. Hsueh, L.-M. Huang, P.-J. Chen, C.-L. Kao, P.-C. Yang, Chronological evolution of IgM, IgA, IgG and neutralisation antibodies after infection with SARS-associated coronavirus, Clinical Microbiology and Infection. 10 (2004) 1062–1066. doi:10.1111/j.1469-0691.2004.01009.x

53. D.N. Ivankov, N.S. Bogatyreva, M.Y. Lobanov, O.V. Galzitskaya, Coupling between Properties of the Protein Shape and the Rate of Protein Folding, PLoS ONE. 4 (2009). doi:10.1371/journal.pone.0006476.

